# TFEB/Mitf links impaired nuclear import to autophagolysosomal dysfunction in C9-ALS

**DOI:** 10.1101/2020.06.26.173021

**Authors:** Kathleen M. Cunningham, Ke Zhang, Kai Ruan, Kirstin Maulding, Mumine Senturk, Jonathan Grima, Hyun Sung, Zhongyuan Zuo, Helen Song, Jeffrey D. Rothstein, Hugo J. Bellen, Thomas E. Lloyd

## Abstract

Disrupted nucleocytoplasmic transport (NCT) has been implicated in neurodegenerative disease pathogenesis; however, the mechanisms by which impaired NCT causes neurodegeneration remain unclear. In a *Drosophila* screen, we identified Ref(2)p/p62, a key regulator of autophagy, as a potent suppressor of neurodegeneration caused by the GGGGCC hexanucleotide repeat expansion (G4C2 HRE) in C9orf72 that causes amyotrophic lateral sclerosis (ALS) and frontotemporal dementia (FTD). We found that p62 is increased and forms ubiquitinated aggregates due to decreased autophagic cargo degradation. Immunofluorescence and electron microscopy of *Drosophila* tissues demonstrate an accumulation of lysosome-like organelles that precedes neurodegeneration. These phenotypes are partially caused by cytoplasmic mislocalization of Mitf/TFEB, a key transcriptional regulator of autophagolysosomal function. Additionally, TFEB is mislocalized and downregulated in human cells expressing GGGGCC repeats and in C9-ALS patient motor cortex. Our data suggest that the C9orf72-HRE impairs Mitf/TFEB nuclear import, thereby disrupting autophagy and exacerbating proteostasis defects in C9-ALS/FTD.

## Introduction

A GGGGCC (G4C2) hexanucleotide repeat expansion (HRE) in chromosome 9 open reading frame 72 (C9orf72) is the most common genetic cause of amyotrophic lateral sclerosis (ALS) and frontotemporal dementia (FTD), accounting for up to 40% of cases of familial ALS (DeJesus-Hernandez et al., 2011; Renton et al., 2011). ALS and/or FTD caused by mutations in C9orf72 (C9-ALS/FTD) is inherited in an autosomal dominant manner, suggesting that the HRE causes disease through gain-of-function or haploinsufficiency (DeJesus-Hernandez et al., 2011; Ling, Polymenidou, & Cleveland, 2013). Loss of C9orf72 function has been linked to disruption of autophagy and lysosome function, though neurodegeneration is not observed in C9orf72 knockout mice (Liu et al., 2016; Y. Shi et al., 2018; Webster et al., 2016), suggesting that C9-ALS/FTD is likely primarily caused by toxicity of the HRE. Furthermore, expression of G4C2 repeats causes neurotoxicity in *Drosophila* and cell culture models of C9-ALS (Goodman et al., 2019; Kramer et al., 2016; Tran et al., 2015). This toxicity has been proposed to occur through either G4C2 repeat RNA-mediated sequestration of RNA-binding proteins or translation of the G4C2 repeats into dipeptide repeat proteins (DPRs) through non-canonical repeat-associated non-AUG translation (Donnelly et al., 2013; Goodman et al., 2019; Mori et al., 2013; Tran et al., 2015).

We previously conducted a *Drosophila* screen of genes that bound with moderate-to-high affinity to G4C2 RNA and identified modulation of the nucleocytoplasmic transport (NCT) pathway as a potent modifier of G4C2 toxicity in both fly and iPS neuron models of C9-ALS (K. Zhang et al., 2015), and this finding has also been made by other groups (Freibaum et al., 2015; Jovicic et al., 2015). The mechanism by which the G4C2 HRE disrupts NCT remain unclear, but potential mechanisms include G4C2 RNA binding to the master NCT regulator RanGAP (K. Zhang et al., 2015), DPRs binding to the nuclear pore complex (Boeynaems et al., 2016; K. Y. Shi et al., 2017; Y. J. Zhang et al., 2016), stress granules sequestering NCT factors (K. Zhang et al., 2018), or through cytoplasmic TDP-43-dependent dysregulation of karyopherin-α (Chou et al., 2018; Gasset-Rosa et al., 2019; Solomon et al., 2018). Recently, a role for NCT disruption in Huntington’s disease and Alzheimer’s disease has been proposed, indicating that NCT disruption may be a common mechanism in many neurodegenerative diseases (Eftekharzadeh et al., 2018; Gasset-Rosa et al., 2017; Grima et al., 2017). However, the pathways affected by NCT disruption that cause neurodegeneration have not yet been elucidated.

In a *Drosophila* screen for modifiers of G4C2-mediated neurodegeneration (K. Zhang et al., 2015), we identified Ref(2)p, the *Drosophila* homolog of p62/SQSTM1 (Sequestosome 1). p62/SQSTM1 is a rare genetic cause of ALS/FTD (Cirulli et al., 2015; Le Ber et al., 2013; Teyssou et al., 2013) and functions in macroautophagy (hereafter termed autophagy) along with other genes implicated in ALS/FTD (Evans & Holzbaur, 2019; Lin, Mao, & Bellen, 2017; Ramesh & Pandey, 2017) such as tank-binding kinase 1 (TBK1), optineurin (OPTN1), ubiquilin 2 and 4 (UBQN2 and 4), valosin-containing protein (VCP), CHMP2B, VapB, and the C9ORF72 protein itself (O’Rourke et al., 2015; Sellier et al., 2016; Sullivan et al., 2016; Ugolino et al., 2016; Webster et al., 2016; Yang et al., 2016). During autophagy, organelles and protein aggregates are degraded via polyubiquitination and targeting to a newly forming autophagosome, followed by degradation upon fusion with the lysosome (Lin et al., 2017).

Although autophagy and nucleocytoplasmic transport have both been implicated in neurodegeneration, it is unclear whether or how these two pathways interact in disease pathogenesis (Gao, Almeida, & Lopez-Gonzalez, 2017; Thomas, Alegre-Abarrategui, & Wade-Martins, 2013). Here, we show that expression of 30 G4C2 repeats is sufficient to disrupt autophagy in *Drosophila*, leading to an accumulation of p62 and ubiquitinated protein aggregates. We find that autophagolysosomal defects are caused by loss of nuclear localization of the transcription factor Mitf (the *Drosophila* homolog of TFEB), which regulates transcription of genes involved in autophagolysosome biogenesis (Bouche et al., 2016; Palmieri et al., 2011; Sardiello et al., 2009; T. Zhang et al., 2015). Furthermore, suppressing this NCT defect is sufficient to rescue Mitf nuclear localization, restoring autophagy and lysosome function and rescuing neurodegeneration. These findings suggest a pathogenic cascade in C9-ALS/FTD whereby NCT disruption causes a failure of autophagosome biogenesis and lysosome dysfunction that ultimately leads to neuronal death.

## Results

### Ref(2)p/p62 knockdown suppresses G4C2-mediated neurodegeneration

Expression of 30 G4C2 repeats (30R) in the eye using *GMR-Gal4* results in progressive photoreceptor degeneration and visible ommatidial disruption by day 15 (Figure 1A)(Xu et al., 2013; K. Zhang et al., 2015). In a genetic modifier screen of over 800 RNAi lines, *UAS-ref(2)p*^*RNAi*^ was among the strongest of 32 suppressors of G4C2-mediated eye degeneration (K. Zhang et al., 2015) (Figure 1A). *ref(2)p* is the *Drosophila* homolog of p62/SQSTM1, and this modifier is of particular interest because *SQSTM1* mutations that cause loss of selective autophagy cause ALS/FTD (Cirulli et al., 2015; Goode et al., 2016; Le Ber et al., 2013), and p62 aggregates are pathological features of both familial and sporadic ALS (Al-Sarraj et al., 2011; Cooper-Knock et al., 2012). Knockdown of *ref(2)p* suppresses eye degeneration, whereas overexpression of *ref(2)p* strongly enhances this phenotype (Figure 1A-B, Figure S1B). Similarly, coexpression of *ref(2)p* RNAi partially rescues the pupal lethality seen with 30R expression in motor neurons (Figure 1C). To determine whether *ref(2)p* knockdown is able to suppress age-dependent neurodegeneration, we used an inducible, pan-neuronal driver (*elavGS-Gal4*) in which 30R-expression leads to a marked reduction in climbing ability after 7 days (Figure 1D). This climbing defect is suppressed with coexpression of *ref(2)p* RNAi suggesting that *ref(2)p* contributes to G4C2-mediated neurotoxicity in the adult brain. *ref(2)p* RNAi coexpression reduces *ref(2)p* levels by about 80% of control and does not alter G4C2 RNA levels (Figure S1C), suggesting that *ref(2)p* acts downstream of G4C2 transcription. Since RAN-translation of arginine-containing DPRs have been implicated in G4C2-mediated toxicity in *Drosophila* (Kwon et al., 2014; Mizielinska & Isaacs, 2014), we next tested whether *ref(2)p* knockdown rescues poly-Glycine-Arginine(GR) repeat-mediated toxicity (Mizielinska & Isaacs, 2014). As shown in Figure S1B, *ref(2)p* RNAi partially rescues the severe eye degeneration phenotype caused by poly(GR)36 expression. Together, these data indicate that *ref(2)p*, the *Drosophila* orthologue of p62/SQSTM1, modulates G4C2-mediated neurodegeneration.

**Figure 1:**
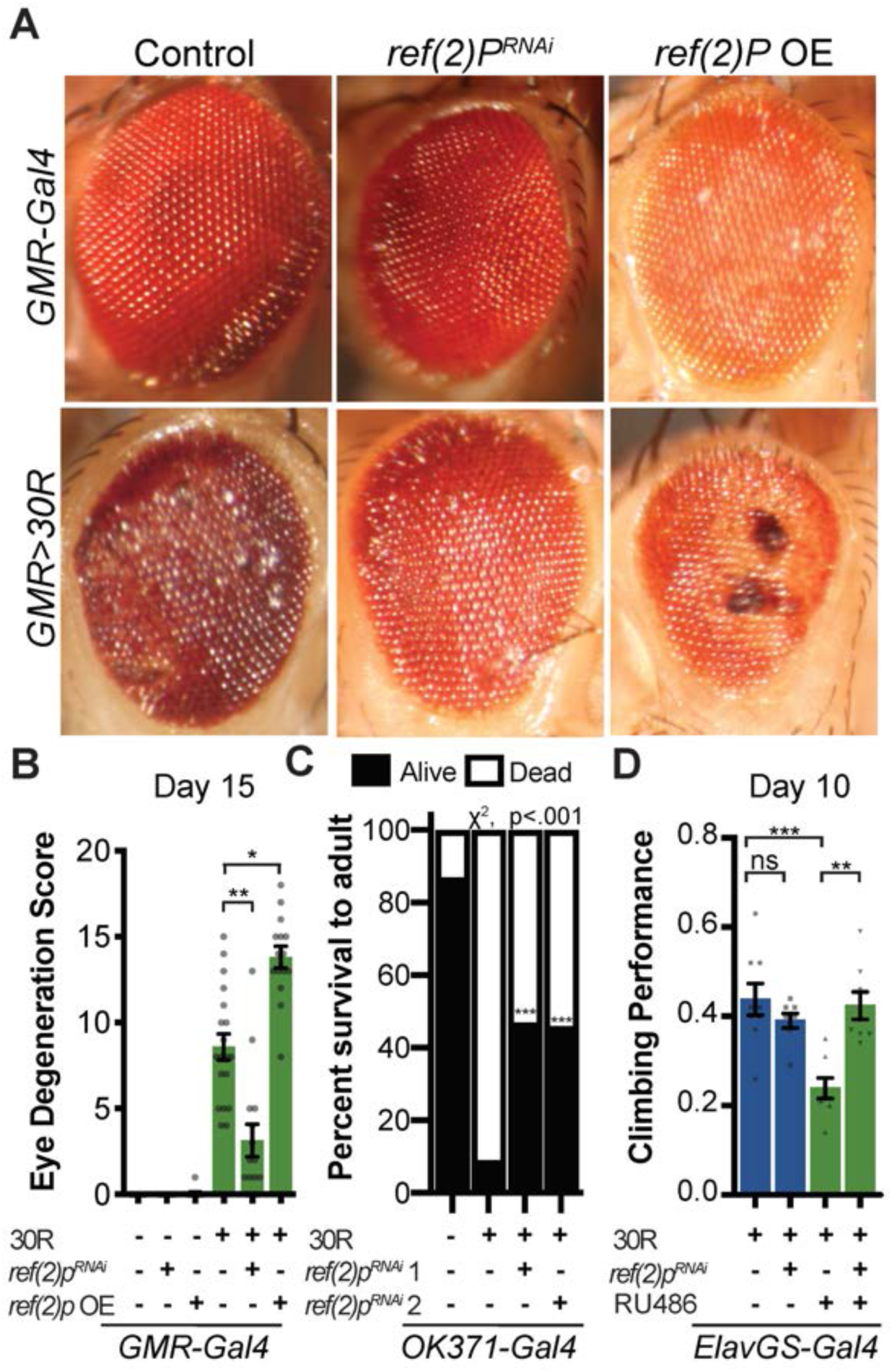
Autophagy receptor Ref(2)p / p62 genetically suppresses 30R-mediated degeneration. (A) 15-day old *Drosophila* eyes expressing *GMR-Gal4* alone or *GMR*>*30R* (bottom row) with *ref(2)p*^*RNAi*^ or overexpression (OE) of HA-tagged *ref(2)p*. (B) Quantification of external eye degeneration, Kruskal-Wallis test, followed by Dunn’s multiple comparisons, n=15. (C) Percent of pupal eclosion in flies expressing 30R under the control of the motor neuron *OK371-Gal4* driver with *OK371-Gal4* alone control, RNAi background control, or Ref(2)p RNAi n≥100 pupae, chi-squared test. (D) Adult flies expressing 30R under the control of the inducible, pan-neuronal elavGS-Gal4 induced with RU486 or vehicle and co-expressing control or *ref(2)p*^*RNAi*^. n= 9, 8, 8, 8 groups of 10 flies. One-way ANOVA with Bonferroni’s Multiple Comparison’s test. Data are mean ± SEM. *:p<.01 **p<.001 ***p<.0001

### G4C2 repeat expression impairs autophagic flux

p62/SQSTM1-positive inclusions are a common pathologic feature seen in brains of C9-ALS/FTD patients where they colocalize with ubiquitin and DPRs (Al-Sarraj et al., 2011). We next investigated the localization of Ref(2)p protein (hereafter referred to as p62) in motor neurons. Expression of 30R leads to the formation of many large p62::GFP puncta in cell bodies compared to controls (Figure 2A-B, Figure S2A) that strongly colocalize with poly-ubiquitinated proteins (Figure 2B). Western blot analysis demonstrates that p62 protein is strongly upregulated in flies ubiquitously expressing 30R (Figure 2C, Figure S2C). Similarly, immunofluorescence staining with a p62 antibody shows endogenous p62 accumulations colocalizing with polyubiquitinated proteins in the ventral nerve cord, eye disc, and salivary gland of flies ubiquitously overexpressing 30R (Figure S2D). These data show that G4C2 repeat expression in fly models recapitulates the p62 protein accumulation with insoluble ubiquitinated protein aggregates seen in C9-ALS/FTD patient tissue and iPS neurons (Almeida et al., 2013; Mackenzie, Frick, & Neumann, 2014).

**Figure 2:**
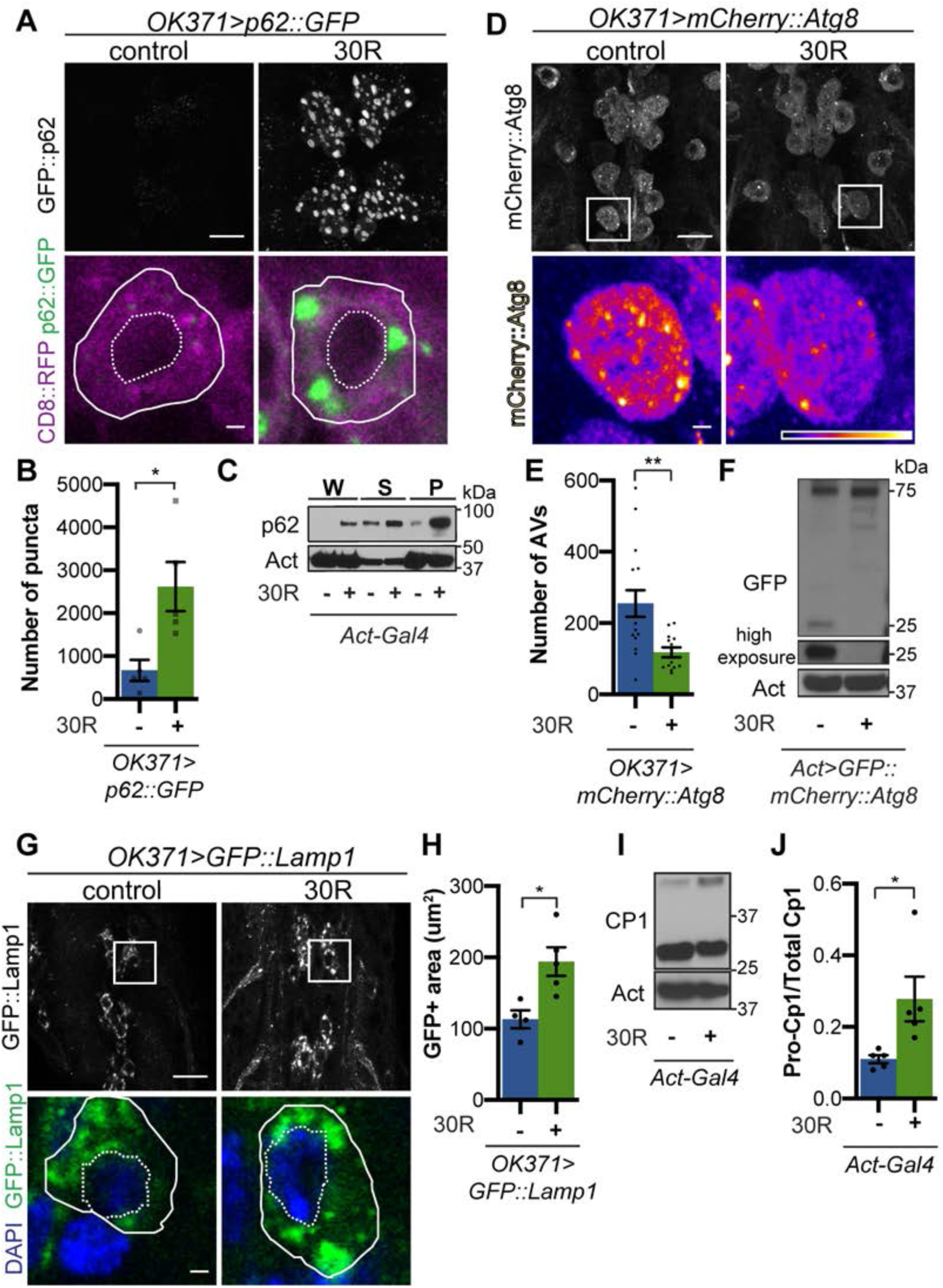
G4C2 repeat expression impairs autophagic flux. (A) *Drosophila* motor neurons expressing *p62::GFP* with or without 30R showing motor neuron cell bodies or a representative single cell body (below). (B) Quantification of p62::GFP puncta in motor neuron cell bodies by number (n= 5 larvae, Mann-Whitney test), (C) Western blot of anti-p62 showing the whole (W), supernatant (S) and pellet (P) fractions of larvae ubiquitously expressing 30R under the control of *Act-Gal4*. (D) Larval motor neurons expressing mCherry::Atg8 with or without 30R showing cell bodies with representative single cell highlighting Atg8-positive puncta (below). (E) Quantification of Atg8-positive autophagic vesicles (AVs) in the ventral nerve cord of *OK371-Gal4*/+ (n=16) or *OK371*>*30R* (n = 13 larvae, Student’s t-test). (F) Anti-GFP Western of whole larvae ubiquitously expressing GFP::mCherry::Atg8 with or without 30R under the control of *Act-Gal4* showing full length GFP::mCherry::Atg8 and cleaved GFP (25 kDa band). G) Larval motor neurons expressing GFP::Lamp (N-terminal (luminal) GFP) with or without 30R in cell bodies. (H) Quantification of GFP::Lamp positive area in (G), n = 5 larvae, Student’s t-test. (I) Western of whole larvae with or without 30R blotted for the lysosomal protease CP1, showing pro-(inactive, top) and cleaved (active, lower) bands. (J) Quantification of the ratio of pro- to cleaved band in (I) (n = 3) *:p<0.05, **p<.001. Data are mean ± SEM.

Increased p62 levels can be due to either increased transcription and/or translation or insufficient protein degradation (Korolchuk, Menzies, & Rubinsztein, 2010). Using qRT-PCR, we show *ref(2)p* transcript levels are unchanged in G4C2 repeat-expressing larvae (Figure S1D), suggesting that G4C2 repeats cause p62 upregulation via inhibiting p62 degradation. Since p62 is degraded by autophagy and, indeed, disrupted autophagic flux is known to cause p62 upregulation, we analyzed autophagy in G4C2-repeat-expressing flies. We first co-expressed the autophagosome marker mCherry::Atg8 (the fly orthologue of mammalian LC3) with 30R in fly motor neurons and found a marked reduction in mCherry::Atg8 vesicles when compared to wild-type controls (Figure 2D-E). p62 accumulation and loss of Atg8 puncta were recapitulated in multiple *Drosophila* models of C9-ALS/FTD (Figure S2E-H). Reduction of Atg8-positive vesicles coupled with the accumulation of p62 and ubiquitin suggest that autophagic flux is impaired in G4C2-expressing animals.

### G4C2 repeat expression causes lysosome defects

To investigate the autophagic pathway defects in G4C2 repeat-expressing neurons, we performed Transmission Electron Microscopy (TEM) on *Drosophila* eyes. As *GMR-Gal4* is expressed throughout the development of photoreceptors (PRs), we chose to perform electroretinograms (ERGs) of fly eyes selectively expressing 30R in photoreceptor neurons using *rh1-Gal4*, which turns on during adulthood. *rh1*>30R PRs show only a mild reduction of ON transient amplitude at 28 days, but complete loss of ON and OFF transients and decrease in ERG amplitude by 56 days (Figure S3A-D), indicating a loss in synaptic transmission and impaired phototransduction pathway respectively. These changes correspond to marked loss of photoreceptors and synaptic terminals by 54 days which are not observed at 28 days (Figure 3A-B; Figure S3E).

**Figure 3.**
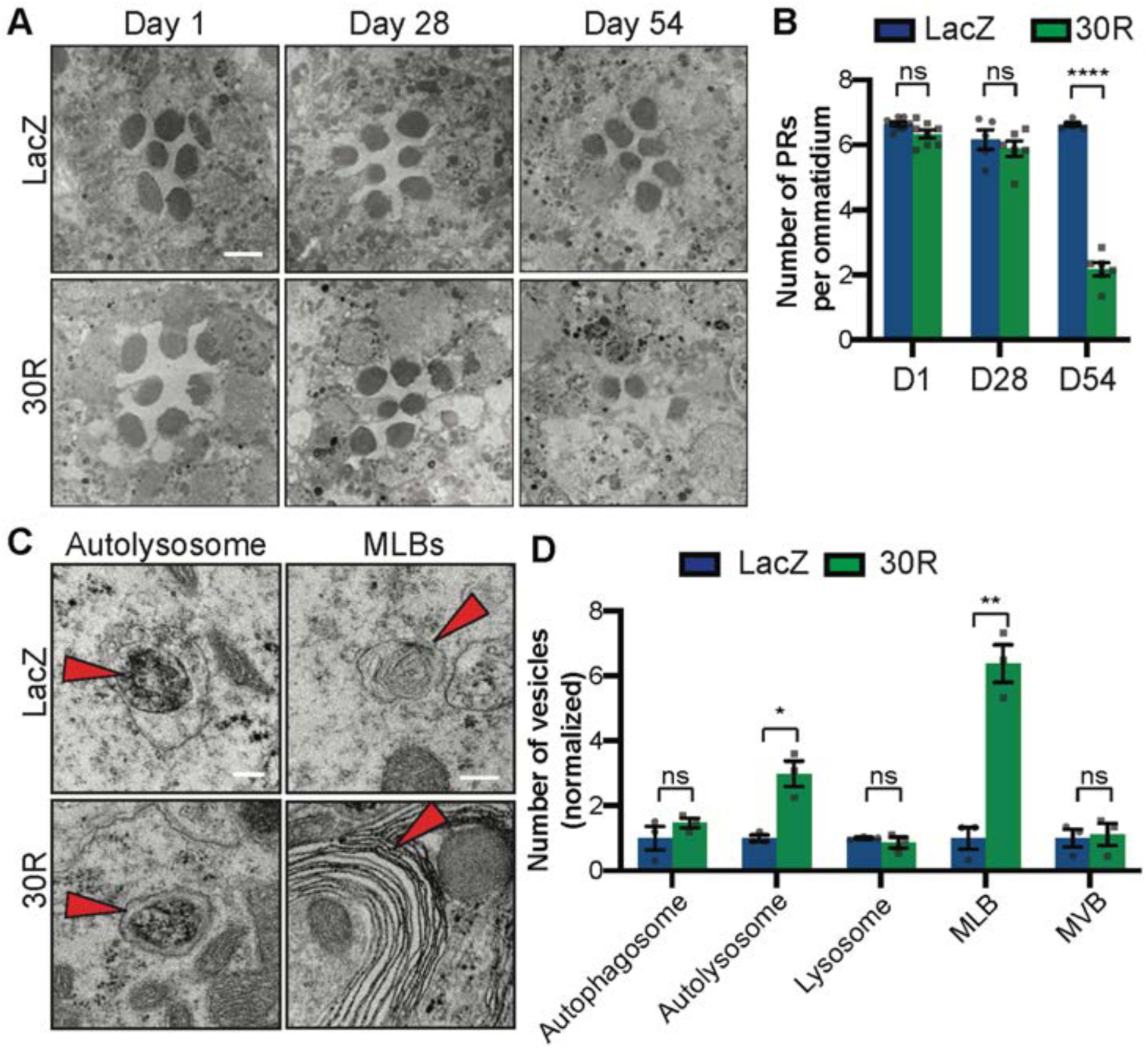
Autophago/lysosomal defects precede neurodegeneration in photoreceptor neurons. (A) TEM of rhabdomeres (cell bodies) in *rhodopsin1-Gal4* driving *LacZ* (control) or 30R at Day 1, Day 28, and Day 54 after eclosion. B) Quantification of number of healthy (not split) photoreceptors per ommatidium in (A), n = 8, 8, 6, 6, 6, 6 flies, Student’s t-test. (C) TEM images at 28 days of *Drosophila* eyes (rhabdomeres) showing representative autolysosomes and multilamellar bodies (MLBs), with or without 30R repeats expressed by *rhodopsin1-Gal4*. D) Quantification to the right of different vesicle types in TEM at day 28 (Student’s t-test, n = 3 adults). Data reported are mean ± SEM. n.s., not significant *, p<0.05, **, p<0.01, ****p<.0001.

We therefore examined autophagic structures by TEM at 28 days, prior to cell loss. Strikingly, we observe a marked increase in the size and number of multilamellar bodies (MLBs) (Figure 3C-D). MLBs are commonly observed in lysosomal storage diseases and result from a deficiency of lysosomal hydrolases and accumulations of lysosomal lipids and membranes (Hariri et al., 2000; Weaver, Na, & Stahlman, 2002). Though we did not detect an alteration in the number of autophagosomes, lysosomes, or multivesicular bodies, we did see a significant increase in the number of autolysosomes (Figure 3C-D). These data suggest that autophagolysosomal function is disrupted in G4C2-expressing photoreceptor neurons at early stages of degeneration.

To further study lysosomal morphology and function, we expressed Lysosome Associated Membrane Protein (LAMP) with luminally-tagged GFP in our control and G4C2-expressing flies. Since GFP is largely quenched by the acidity of lysosomes in control animals (Pulipparacharuvil et al., 2005), the accumulation of GFP::Lamp-positive vesicles in 30R-expressing motor neurons suggests a defect in lysosomal acidity or targeting of GFP::Lamp to mature lysosomes (Figure 2G-H). Furthermore, we observe a marked increase in size and number of late endosomes and lysosomes using genomically tagged Rab7::YFP throughout 30R-expressing motor neurons (Figure S4A) without alterations in early endosomes labeled with Rab5:YFP (data not shown). Together, these data demonstrate a marked expansion of the late endosome/lysosome compartment in G4C2-expressing neurons.

Though accumulation of p62 and ubiquitinated proteins could be caused by a failure of autophagic vesicles to fuse with the degradative endolysosomal compartment, we did not detect a decrease in Atg8+, Rab7+ amphisomes in G4C2-expressing motor neuron cell bodies (Figure S4B-F). To assess autophagolysosomal function after fusion, we performed a “GFP liberation assay” on larvae expressing GFP::mCherry::Atg8 (Klionsky et al., 2016; Mauvezin, Ayala, Braden, Kim, & Neufeld, 2014). GFP is degraded more slowly than the rest of the Atg8 protein, leaving a population of free GFP in functioning lysosomes. Free lysosomal GFP is not observed in G4C2-expressing larvae, suggesting an impairment in GFP::mCherry::Atg8 degradation by the lysosome (Figure 2F). To directly probe lysosome enzymatic activity, we performed Western analysis of *Drosophila* cathepsin CP1. Whereas pro-CP1 is normally cleaved to its mature form by acid hydrolases in lysosomes (Kinser & Dolph, 2012), larvae ubiquitously expressing 30R show an increase in the ratio of pro-CP1 to CP1, indicating a decrease in pro-CP1 cleavage efficiency (Fig 2I-J). Together, these data suggest that lysosomes are expanded and dysfunctional in G4C2 repeat-expressing animals.

Given the impairment in autophagic flux, we hypothesized that genetic or pharmacologic manipulations that accelerate autophagy may suppress neurodegenerative phenotypes, whereas those that further impede autophagy would enhance the phenotypes. Indeed, in a candidate-based screen, activation of early steps in the autophagic pathway (e.g. by Atg1 overexpression) suppresses eye degeneration and blocking autophagosome/lysosome fusion (e.g. by Snap29 knockdown) enhances eye degeneration (Table S1). Similarly, pharmacologic activation of autophagy via inhibition of mTOR with rapamycin or mTor-independent activation via trehalose rescues neurodegenerative phenotypes and p62 accumulation (Figure S4G-J). Together, these data show that promoting autophagy or lysosomal fusion are potent suppressors of G4C2-mediated neurodegeneration.

### Nucleocytoplasmic transport impairment disrupts autophagic flux

A diverse array of cellular pathways including autophagy, RNA homeostasis, and nucleocytoplasmic transport are implicated in the pathogenesis of ALS and FTD (Balendra & Isaacs, 2018; Evans & Holzbaur, 2019; Gao et al., 2017; Lin et al., 2017; Ling et al., 2013; Ramesh & Pandey, 2017). However, the sequence of events in the pathogenic cascade remains unknown. Cytoplasmic protein aggregates or RNA stress granule formation is sufficient to disrupt nucleocytoplasmic transport (Woerner et al., 2016; K. Zhang et al., 2018). We therefore tested whether defects in autophagy are upstream, downstream, or in parallel with defects in nucleocytoplasmic transport.

We first tested whether knockdown of *ref(2)p* rescues the mislocalization of the NCT reporter shuttle-GFP (S-GFP) containing both a nuclear localization sequence (NLS) and nuclear export sequence (NES). G4C2 repeat expression causes mislocalization of S-GFP to the cytoplasm (K. Zhang et al., 2015), but knockdown of *ref(2)p* does not restore nuclear localization (Figure S5A). Similarly, stimulation of autophagy with rapamycin or trehalose fails to rescue S-GFP mislocalization in G4C2 expressing salivary glands (Figure S5B). Restoring autophagy does not rescue NCT defects although it can rescue neurodegeneration, suggesting that autophagy defects are either independent of or downstream of NCT defects. Indeed, *RanGAP* knockdown is sufficient to increase the number and size of p62::GFP puncta, similar to the effects of overexpressing the G4C2 repeats (Figure S5C), suggesting that NCT disruption is sufficient to disrupt autophagic flux in *Drosophila* motor neurons.

### Mitf is mislocalized and inactivated in Drosophila models of C9-ALS/FTD

Because we observed a reduction in autophagosomes and expansion of lysosome-related organelles, we hypothesized that transcription factors regulating autophagolysosomal function may be mislocalized to the cytoplasm due to disrupted nuclear import. The TFE family of transcription factors (TFEB, TFE3, MITF, and TFEC) regulates multiple steps of autophagy from autophagosome biogenesis through lysosome acidification via a network of genes called the Coordinated Lysosome Expression And Regulation (CLEAR) network (Settembre et al., 2011). These transcription factors are regulated by phosphorylation of canonical nuclear export (NES) and nuclear import (NLS) sequences (Li et al., 2018). In *Drosophila*, this conserved transcription factor family is represented by a single homolog called Mitf (Bouche et al., 2016; T. Zhang et al., 2015).

Mitf knockdown in the nervous system causes lysosomal defects similar to those we observe in G4C2-expressing flies (Bouche et al., 2016; Hallsson et al., 2004; Sardiello et al., 2009; Song et al., 2013). Additionally, TFEB levels are reduced in SOD1 cell culture and mouse ALS models (Chen et al., 2015) as well as in ALS and Alzheimer’s patient brain tissue (Wang, Wang, Xu, & Lakshmana, 2016). Therefore, we hypothesized that impaired Mitf nuclear import might underlie the autophagolysosomal phenotypes in fly models of C9-ALS. Indeed, both salivary gland cells and motor neurons expressing 30R have a marked reduction in nuclear to cytoplasmic ratio of Mitf (Figure 4A-D). To assess whether this loss of nuclear Mitf leads to a decrease in CLEAR gene expression in adult heads we expressed 30R using a ubiquitous inducible driver, *daughterless-GeneSwitch (daGS)*. In control flies, a mild (∼1.75 fold) overexpression of *Mitf* mRNA resulted in a significant upregulation of 3 of the 7 Mitf targets tested (the v-ATPase subunits *Vha16-1, Vha68-2, and Vha44*) and upregulation of 4 others (Figure 4E), although not statistically significant. Importantly, coexpression of 30R with *daGS*>*Mitf*; led to a similar ∼2x increase in *Mitf* transcripts but did not induce Mitf target genes (Figure 4E). This lack of Mitf target induction in 30R flies suggests that decreased nuclear import of Mitf suppresses the ability of 30R-expressing flies to upregulate CLEAR genes in order to maintain autophagic flux.

**Figure 4.**
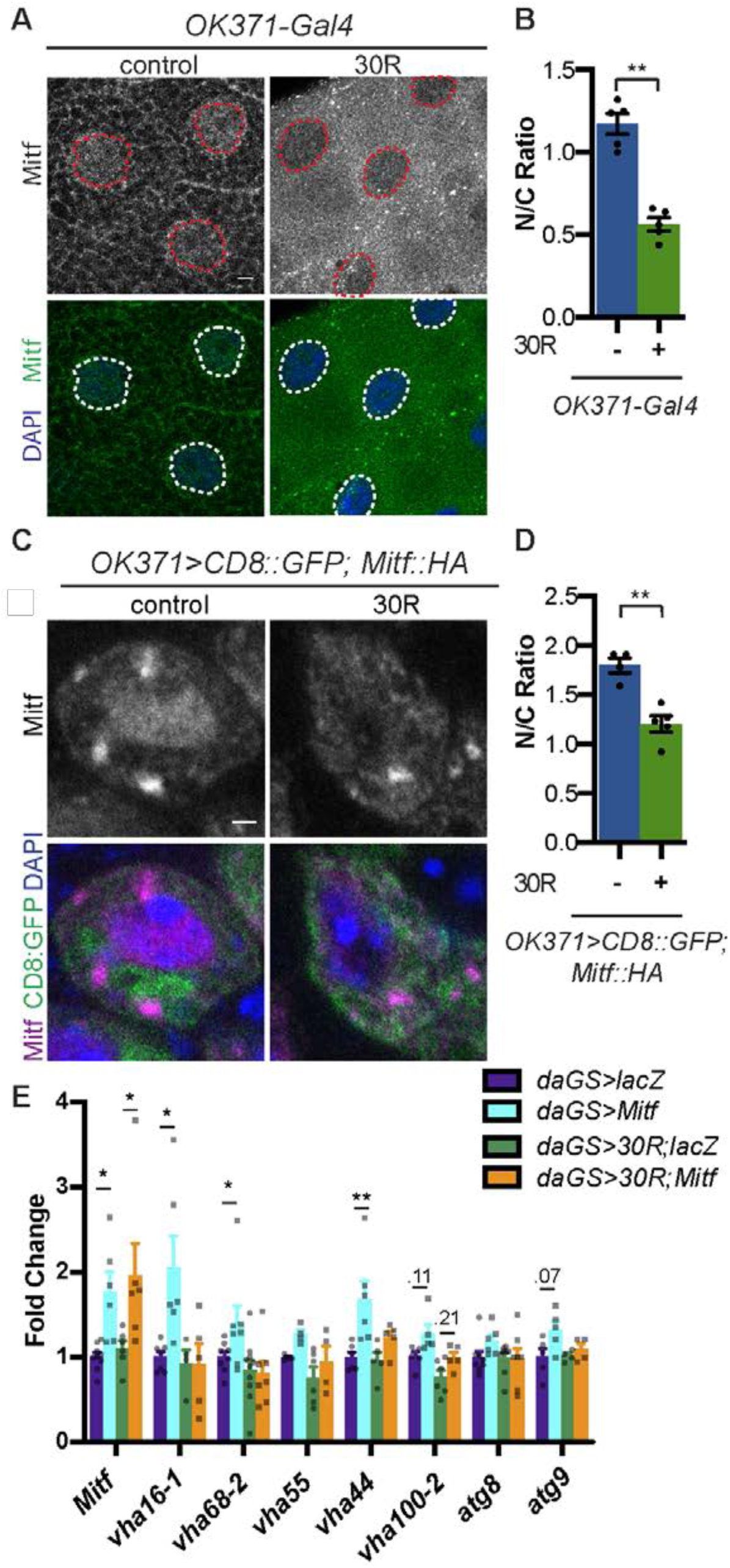
Mitf/TFEB is mislocalized from the nucleus and inactivated. (A) Larval salivary glands stained with anti-Mitf and DAPI with or without 30R expression. (B) Quantification of the nuclear to cytoplasmic (N/C) ratio of Mitf intensity (Student’s t-test, n = 5 larvae). (C) *Drosophila* motor neurons (MNs) expressing *UAS-Mitf-HA* with either control or 30R stained with anti-Mitf and DAPI to show nuclear localization. (D) Quantification of nuclear to cytoplasmic (N/C) ratio of Mitf intensity, (Student’s t-test, n = 5 larvae, ≥ 10 MNs/VNC). (E) Quantitative RT-PCR of the levels of Mitf transcripts and 7 target genes in *Drosophila* heads expressing control (*UAS-lacZ*) or 30R driven by *daGS-Gal4* in control conditions or with overexpression of Mitf (Biological replicates of 30 heads, One way ANOVA with Sidak’s multiple comparisons, n ≥ 4). Data reported are mean ± SEM. n.s., not significant *, p<0.05, **, p<0.01.

We next examined whether rescue of nucleocytoplasmic transport defects in 30R-expressing animals can rescue Mitf nuclear localization and autophagolysosomal defects. Mitf/TFEB nuclear export has recently been demonstrated to be regulated by exportin-1 (Li et al., 2018; Silvestrini et al., 2018). Knockdown of exportin-1 (*Drosophila emb*) rescues G4C2-mediated cytoplasmic Mitf mislocalization in the salivary gland (Figure 5A-B) and GFP::Lamp accumulation in motor neurons (Figure 5C-D). Importantly, *emb* knockdown increases the total number of autophagosomes in G4C2-expressing motor neuron cell bodies by ∼3-fold (Figure 5E-F), suggesting that nuclear retention of Mitf rescues autophagolysosomal defects. Together, these data demonstrate that autophagolysosomal dysfunction in 30R-expressing animals occurs downstream of nucleocytoplasmic transport disruption.

**Figure 5.**
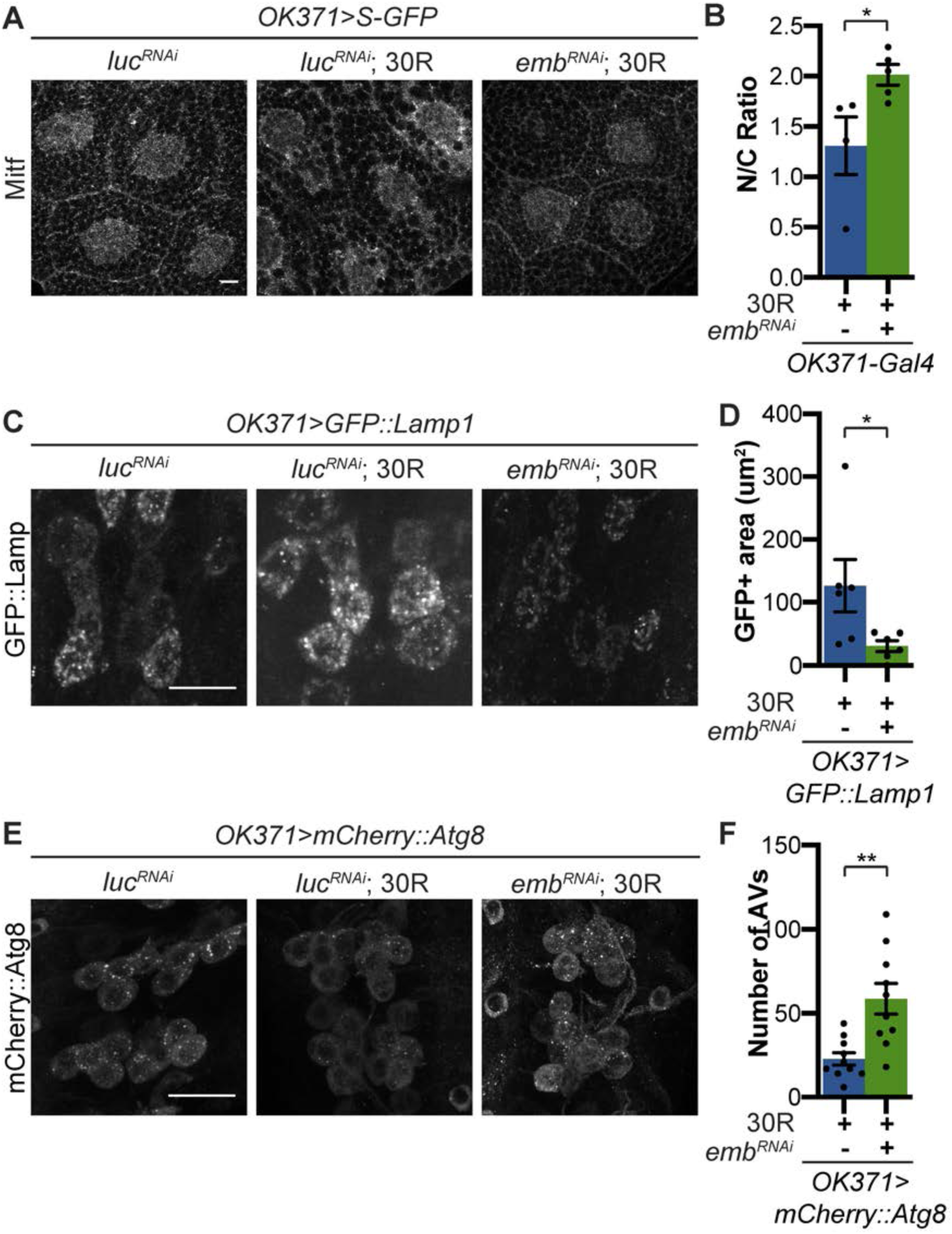
Modulation of nucleocytoplasmic transport rescues autophagy and lysosome dysfunction. A) Larval salivary glands stained with anti-Mitf and DAPI with or without 30R, expressing either control or *emb*^*RNAi*^ exportin) and shuttle-GFP (S-GFP). B) Quantification of Mitf N/C ratio in (A), (Student’s t-test, n= 5 larvae). C) *Drosophila* motor neurons expressing *UAS-Lamp::GFP* with N-terminal (luminal) GFP with 30R and either control (*luciferase (luc) RNAi*) or *emb*^*RNAi*^. D) Quantification of C (Student’s t-test, n = 6 larvae). E) Larval motor neurons expressing UAS-mcherry::Atg8 with 30R and either control (luciferase RNAi) or embRNAi. F) Quantification of (E), Student’s t-test, n = 10 larvae. Data reported are mean ± SEM. *, p<0.05, **, p<0.01.

### Mitf rescues G4C2 repeat-mediated degeneration

Since Mitf mislocalization contributes to autophagolysosome defects in a fly C9-ALS model, we hypothesized that increasing total levels of Mitf might compensate for impaired nuclear import. While high level *Mitf* overexpression is toxic in *Drosophila* (Hallsson et al., 2004), a genomic duplication of the *Mitf* gene (*Mitf Dp*) is sufficient to partially rescue 30R-mediated eye degeneration, while *Mitf* knockdown strongly enhances eye degeneration (Figure 6A-B). Furthermore, pupal lethality caused by 30R expression in motor neurons and climbing impairment in *elavGS*>30R flies are also partially rescued by *Mitf Dp* (Figure 6C-D). Therefore, increasing Mitf levels in multiple neuronal subtypes in *Drosophila* suppresses G4C2-mediated neurotoxicity, consistent with our hypothesis that loss of nuclear Mitf is a key contributor to G4C2-mediated neurodegeneration.

**Figure 6.**
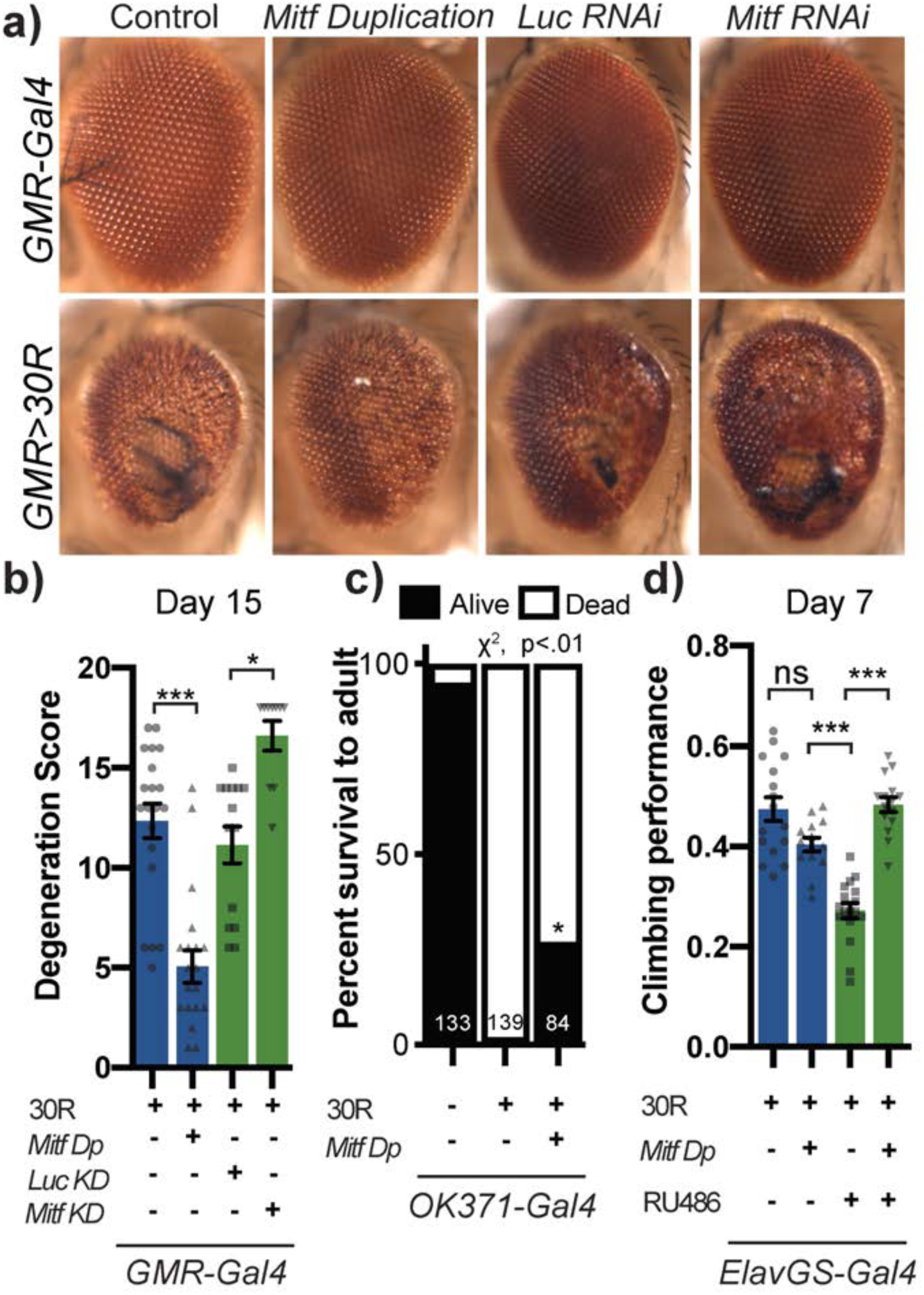
Transcription factor Mitf/TFEB suppresses neurodegeneration caused by G4C2 expansion. (A) 15-day old fly eyes expressing 30R under the control of *GMR-Gal4*, crossed to either control, genomic *Mitf Duplication* (*Mitf Dp*), Luciferase (Luc) RNAi as control, or *Mitf*^*RNAi*^. (B) Quantification of external eye degeneration shown in (A), Kruskal-Wallis test followed by Dunn’s multiple comparisons, n= 10-20. (C) Percent of pupal eclosion in flies expressing 30R under control of *OK371-Gal4* driver with *Gal4* alone control, background control, or *Mitf Dp*, Chi-square test, n= 139, 84 respectively. (D) Adults expressing 30R under control of the inducible, pan-neuronal *elavGS-Gal4* driver have decreased climbing ability at 7 days age. Co-expressing *Mitf Dp* with the 30R rather than control *UAS-lacZ* rescues climbing ability. Data are mean ± SEM. n.s., not significant *, p<0.05, ***p<0.001

If the impaired lysosomal function we observe in our *Drosophila* model is contributing to neurodegeneration downstream of NCT defects, we would predict that genetic upregulation of key regulators of lysosome function may suppress degenerative phenotypes. Indeed, overexpression of *rab7*, the small GTPase required for fusion of autophagosomes with lysosomes, or *TRPML*, a lysosomal calcium channel, suppress eye degeneration (Figure S6A-B). Furthermore, overexpression of key lysosomal v-ATPase subunits whose expression is regulated by *Mitf* also suppresses neurodegeneration in the *Drosophila* eye, while RNAi-mediated knockdown enhances degeneration (Figure S6A-B). Interestingly, loss of the ALS-associated gene *ubqn* in *Drosophila* was also rescued by increase in key lysosomal v-ATPases or by nanoparticle mediated lysosome acidification (Senturk et al., 2019). Overexpression of these Mitf-regulated genes also showed partial rescue of pupal lethality in animals expressing 30R in motor neurons (Figure S6C). These findings suggest a model whereby downregulation of Mitf targets leads to lysosomal disruption in G4C2 repeat-expressing flies.

### Nuclear TFEB is reduced in human cells and motor cortex with GGGGCC repeat expansions

In humans, TFEB is the homolog of *dMitf* that is best characterized for its role in autophagy and has been implicated in neurodegenerative disease (Cortes & La Spada, 2019; Martini-Stoica, Xu, Ballabio, & Zheng, 2016). Interestingly, nuclear TFEB was selectively depleted in the motor cortex of a sample of 5 ALS patients compared to 5 controls (Wang et al., 2016). To test the relevance of our findings in *Drosophila* models to human disease, we next examined whether G4C2 repeat expression impairs nuclear import of TFEB in HeLa cells stably expressing TFEB-GFP (Roczniak-Ferguson et al., 2012) using a 47-repeat (47R) G4C2 construct that expresses tagged DPRs (See methods). TFEB-GFP is predominantly localized to the cytoplasm in both control cells and cells expressing a 47 G4C2 repeat construct (Figure 7A-B). However, induction of autophagy by 3 hours starvation leads to nuclear translocation of TFEB in control cells but not in cells expressing 47R (Figure 7A-B). We then tested the effect of expression of DPRs produced by alternate codons (i.e. in the absence of G4C2 repeats): poly-(GA)50, poly-(GR)50, and poly-(PR)50 (Figure S7A-B). Only polyGA causes a mild impairment in TFEB nuclear translocation, whereas polyGR and polyPR stimulate TFEB-GFP nuclear translocation even under normal culture conditions. These data suggest that despite the protein stress response initiated by DPRs driving nuclear translocation of TFEB under homeostatic conditions, human cells expressing the G4C2 repeat are unable to efficiently import TFEB into the nucleus.

**Figure 7.**
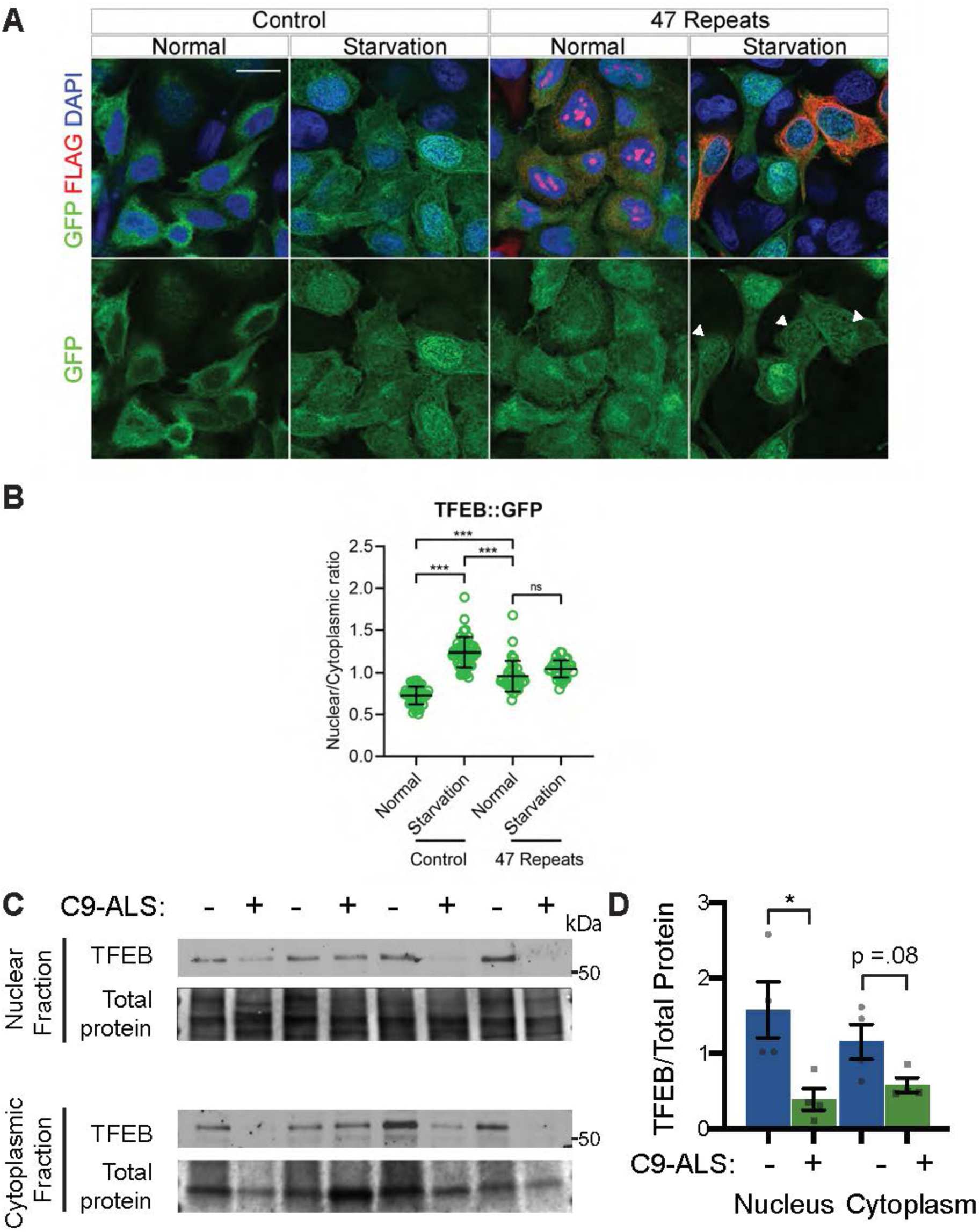
Nuclear TFEB is reduced in human cells expressing GGGGCC and C9-ALS human motor cortex. (A) HeLa cells stably expressing TFEB::GFP transfected with 0R (Control) or a 47R construct in normal (DMEM) or starved (3 hour EBSS) conditions. White arrows indicate transfected cells in the 47R starved group. (B) Quantification of cells from (A) showing the nucleocytoplasmic ratio of TFEB for each group (n = n = 47, 47, 35, 38 cells, Kruskal-Wallis test with Dunn’s multiple comparisons). (C) Western blot for TFEB of human motor cortex samples fractionated into cytoplasmic and nuclear samples from non-neurological controls and C9-ALS patients. (B) Quantification of TFEB levels against total protein loading (Faststain) in non-neurological controls and C9-ALS patients (N = 4, Student’s t-test). Data reported are mean ± SEM. n.s., not significant *, p<0.05, ***p<0.001.

To further investigate the relevance of loss of TFEB nuclear import to C9-ALS patients, we obtained human motor cortex samples from 4 non-neurological controls and 4 C9-ALS patients (Table S2). These samples were fractionated into cytoplasmic and nuclear-enriched fractions and assayed for TFEB using Western analysis. TFEB is reduced by an average of 76% in the nuclear fraction and by about 50% in the cytoplasm in C9-ALS compared to controls (Figure 7C-D, Figure S7C). These data suggest that TFEB protein is downregulated in C9-ALS/FTD motor cortex, but the greatest depletion occurs in the nucleus. Therefore, we propose a model whereby disruption of protein nuclear import by the *C9orf72*-HRE results in a failure of Mitf/TFEB to translocate to the nucleus to regulate the autophagic response to protein stress. This impairment in autophagic flux results in the accumulation of p62-positive ubiquitinated aggregates that contribute to a feed-forward cycle of proteostasis disruption and chronic cell stress signaling, eventually leading to neuron cell death.

## Discussion

Our work has revealed that the ALS-associated G4C2 hexanucleotide repeat is sufficient to disrupt multiple aspects of autophagy. In *Drosophila*, G4C2 repeats cause loss of autophagosomes and disrupt lysosomal structure and function. This accumulation of autolysosomes and lysosome-related organelles called multilamellar bodies (MLBs) has been observed in lysosomal storage disorders and has been reported in spinal cord tissue from sporadic ALS patients (Bharadwaj, Cunningham, Zhang, & Lloyd, 2016; Parkinson-Lawrence et al., 2010; Sasaki, 2011). Regulation of protein and lipid homeostasis by the lysosome may be particularly important in neurons since they are post-mitotic and have high energy demands (Fraldi, Klein, Medina, & Settembre, 2016). Loss of function of *C9ORF72* also disrupts autophagy and lysosomal function in multiple cell types (Farg et al., 2014; Ji, Ugolino, Brady, Hamacher-Brady, & Wang, 2017; O’Rourke et al., 2015; Sellier et al., 2016; Y. Shi et al., 2018; Sullivan et al., 2016; Ugolino et al., 2016; Webster et al., 2016; Yang et al., 2016; Zhu et al., 2020), suggesting a mechanism whereby G4C2 repeats may have synergistically detrimental effects with haploinsufficient *C9ORF72* in C9-ALS/FTD patients. Additionally, multiple forms of familial ALS are caused by genes that regulate autophagy and lysosome function (Evans & Holzbaur, 2019; Lin et al., 2017; Ramesh & Pandey, 2017), and upregulation of lysosome function has been proposed to be beneficial in multiple preclinical models of ALS (Donde et al., 2020; Mao et al., 2019; Senturk et al., 2019; Y. Shi et al., 2018). Thus, our findings suggest that like other forms of ALS, neurotoxicity of G4C2 repeats in C9 ALS-FTD is at least partially caused by disrupted autophagolysosomal function.

The finding that *ref(2)p* knockdown prevents or delays G4C2-mediated neurodegeneration is surprising, as p62/SQSTM1 is thought to link toxic ubiquitinated aggregates to LC3 to remove aggregates via selective autophagy (Cipolat Mis, Brajkovic, Frattini, Di Fonzo, & Corti, 2016; Levine & Kroemer, 2008; Saitoh et al., 2015). However, other studies have also suggested that p62 may contribute to (rather than ameliorate) toxicity of ubiquitinated protein aggregates. For example, *Atg7*^*-/-*^ mice display severe defects in autophagy and accumulation of p62-positive protein aggregates in the liver and brain, and knockout of p62 in these mice prevents the formation of ubiquitinated aggregates and rescues liver dysfunction via suppression of chronic oxidative stress signaling (Komatsu et al., 2007). Additionally, ATM-mediated DNA double stranded break repair is impaired in cultured neurons expressing the *C9orf72*-HRE, and this phenotype is rescued by p62 knockdown (Walker et al., 2017). These findings suggest that accumulation of p62-positive aggregates may contribute to DNA damage previously described in C9-ALS. Further, p62 is found to co-localize with DPRs in C9-ALS patients (Al-Sarraj et al., 2011; Mackenzie et al., 2014; Mori et al., 2013) and may promote protein aggregation. We hypothesize that p62-positive aggregate formation in C9-patients contributes to neurotoxicity by activating downstream signaling pathways that are alleviated by autophagy-mediated aggregate clearance.

While many groups have reported nucleocytoplasmic transport dysfunction in ALS, it has remained unclear how NCT disruption causes ALS. Stress granules can recruit nuclear pore proteins to the cytoplasm and cause nucleocytoplasmic transport defects, suggesting that the disruptions in phase separation of RNA binding proteins may lie upstream of nucleocytoplasmic transport defects (K. Zhang et al., 2018). Recently, Ortega et al. discovered that hyperactivity of nonsense-mediated decay (NMD) may lie downstream of nucleocytoplasmic transport, indicating that multiple proteostasis pathways may be disrupted (Ortega et al., 2020). Additionally, selective autophagy has been proposed to be required for nuclear pore turnover (Lee et al., 2020), implying that autophagy defects may contribute to the cytoplasmic nuclear pore pathology found in C9-ALS patients and animal models. Our data show that in *Drosophila*, HeLa cells, and C9-ALS postmortem brain tissue, nucleocytoplasmic transport defects lead to an inability to activate TFEB translocation to the nucleus, causing widespread autophagy defects and accumulation of protein aggregates (**Figure 8**). These findings place nucleocytoplasmic transport defects in ALS upstream of proteostasis defects.

**Figure 8.**
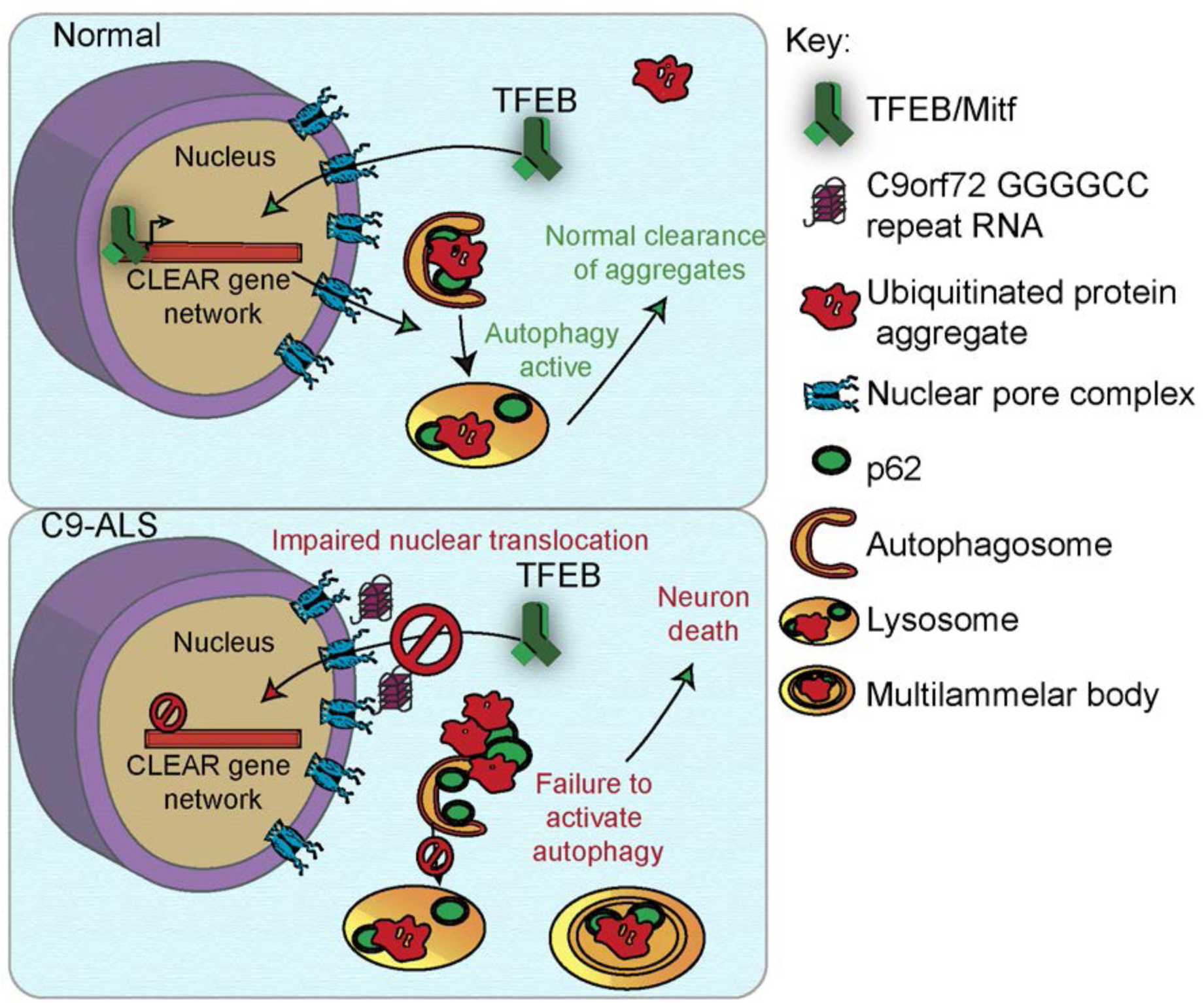
A proposed model of autophagolysosomal dysfunction in C9-ALS. G4C2 repeat expansion causes nucleocytoplasmic transport disruption through multiple proposed mechanisms including G4C2 RNA binding of RanGAP and stress granule recruitment of nucleocytoplasmic transport machinery. Transport disruption impairs nuclear translocation of autophagy-mediating transcription factors such as Mitf/TFEB in response to proteostatic stress. Failure to induce autophagic flux leads to accumulation of ubiquitinated protein aggregates colocalized with Ref(2)p/p62 and large, non-degradative lysosomes and MLBs. Ref(2)p/p62 accumulation contributes to chronic protein stress signaling and eventually neuronal cell death.

Importantly, TFEB has been previously proposed as a therapeutic target in ALS and other neurodegenerative disease (Cortes & La Spada, 2019). Upregulation of TFEB signaling helps clear multiple types of proteotoxic aggregates found in Alzheimer’s disease, Parkinson’s disease, Huntington’s disease, ALS and FTD (Decressac et al., 2013; Parr et al., 2012; Polito et al., 2014; Torra et al., 2018; Vodicka et al., 2016). Our study suggests that modulation of TFEB nucleocytoplasmic transport may be an additional therapeutic target, and that targeting both nucleocytoplasmic transport and autophagy may act synergistically in ALS and FTD.

## Materials and Methods

### Drosophila genetics

A complete list of fly lines used in this study is described in Supplemental Table 3. *Drosophila* were raised on standard cornmeal-molasses food at 25°C. For eye degeneration, *GMR-GAL4, UAS-30R*/CyO, *twi-GAL4, UAS-GFP* were crossed to *UAS-modifier* lines or background controls and *GMR-GAL4, UAS-30R*/*UAS-modifier* or *GMR-GAL4, UAS-30R/*+ were selected (where *UAS-modifier* can be on any chromosome) from the offspring and aged at 25°C for 15 days. Eye degeneration is quantified using a previously described method^89^. Briefly, points were added if there was complete loss of interommatidial bristles, necrotic patches, retinal collapse, loss of ommatidial structure, and/or depigmentation of the eye. Eye images were obtained using a Nikon SMZ 1500 Microscope and Infinity 3 Luminera Camera with Image Pro Insight 9.1 software.

For pupal survival assay, either 3 males from *OK371-Gal4* or *OK371-Gal4; UAS-30R*/TM6G80(Tb) were crossed to 5-6 female flies containing UAS-modifier lines or background controls. Parental adult crosses were transferred to fresh vials every 2-3 days. After 15 days, non-tubby pupated flies that were (either *OK371-Gal4/UAS-modifier, OK371*/+; *UAS-30R*, or *OK371-Gal4*/*UAS-modifier*; *UAS-30R* were scored as either eclosed (empty pupal case) or non-eclosed (typically a fully developed pharate adult fly unable to eclose from pupal case due to paralysis).

For the climbing assay, *UAS-30R*; *elavGS-Gal4* were crossed to experimental or genetic background controls. Adults were transferred 3-5 days after eclosion to vials containing 200 μM RU486 food or ethanol vehicle alone and transferred to new vials every 2-3 days. After aging 7-10 days, groups of 10 flies were placed into empty food vials and were tapped to the bottom and then locomotor function assessed by their negative geotaxis (flies reflexively crawl against gravity) response as measured by ability to climb 8 cm in 10 seconds. Each cohort of 10 flies was tested 10 times to obtain an average. N represents individual cohorts of 10 flies.

### *Drosophila* drug feeding

Cornmeal-molasses-yeast fly food was melted and then cooled for 5 minutes before being mixed with concentrations of mifepristone (RU486, Sigma), rapamycin (Selleckchem), or trehalose (Sigma) and cooled to room temperature. Ethanol or DMSO was used as a vehicle control. Parent flies were crossed on normal food, and then they were transferred to food containing drug every 2-3 days such that their offspring would develop in food containing drug or adult offspring were transferred to drug food once eclosed as noted. Wandering third instars were selected for immunostaining or western blot analysis. Adult flies were aged on the drug-containing food for 15 days before analyzing their eye morphology or assessed for climbing ability on the day noted.

### Quantitative RT–PCR

For each genotype, mRNA was collected from 5 flies using the TRIzol reagent (Thermo Fischer Scientific) following the manufacturer’s protocol. Reverse transcription was performed using SuperScript III First-Strand synthesis kit (Thermo Fischer Scientific) following the manufacturer’s protocol. Quantitative PCR was performed using SYBR Green PCR system (Applied Biosystem) on a 7900HT fast Real-Time PCR system (Applied Biosystem). Primers are listed in Supplemental Table 4. The primers for G4C2 repeats were designed to amplify a 3’ region immediately after the repeats in the UAS construct.

### Immunofluorescence staining and imaging

For *Drosophila* ventral nerve cords, wandering 3^rd^ instar larvae were dissected in HL3^90^ using a standard larval fillet dissection then fixed in 4% paraformaldehyde (or, for *UAS-Atg8-Cherry* experiments Bouin’s fixative (Sigma)) for 20 minutes, followed by wash and penetration with PBS 0.1% Triton X-100 (PBX) for 3x 20-minute washes. The tissues were blocked for 1 hour at room temperature in PBS with 5% normal goat serum (NGS) and 0.1% PBX, then stained with the following primary antibodies at 4C overnight (16h): rabbit anti-dsRed (Clontech) 1:1,000; mouse anti-poly ubiquitin FK1 (Enzo Life Sciences) 1:200; rabbit anti-Ref2p (a gift from Gabor Juhasz, Eotvos Lorand University, Budapest, Hungary]; guinea pig anti-Mitf [a gift from Francesca Pignoni, SUNY Upstate, New York, USA) 1:500; rat anti-HA (Sigma) 1:200. Tissues were washed 3 times for 20 minutes each with 0.1% PBX. Secondary antibodies (Goat antibodies conjugated to Alexa Fluor 568, 488, 633) diluted in 0.1% PBX 5% NGS and incubated for 2h and then washed 3 times for 20 minutes each with 0.1% PBX. During one wash, DAPI was added to the prep at a final concentration of 1ug/mL. Larvae were mounted in Fluoromount-G (Invitrogen).

*Drosophila* salivary glands were dissected using a standard protocol and stained as above excepting for stronger solubilization with 0.3% PBX. Fixed cells or tissues were analyzed under an LSM780 or LSM800 confocal microscope (Carl Zeiss) with their accompanying software using Plan Apochromat 63 ×, NA 1.4 DIC or Plan Apochromat 40×, 1.3 Oil DIC objectives (Carl Zeiss) at room temperature. Images were captured by an AxioCam HRc camera (Carl Zeiss) and were processed using ImageJ/Fiji (National Institutes of Health). To quantify fluorescent intensities, after opening the images in ImageJ/Fiji, certain areas/bands were circled and the intensities were measured. Puncta were counted using the Analyze Particles function in Image J using the same thresholding across experiments. Images are representative and experiments were repeated two to five times.

### Western Blotting

Tissues or cells were homogenized and/or lysed in RIPA buffer (50 mM Tris-HCl pH 7.4, 150 mM NaCl, 0.1% SDS, 0.5% sodium deoxycholate, and 1% Triton X-100) supplemented with protease inhibitor cocktail (Complete, Roche) using microcentrifuge pestles, and then were incubated in RIPA buffer on ice for 20 minutes. Samples were spun down at 100g for 5 minutes to remove carcass and unbroken cells. For protein quantification, solution was diluted and measured by BCA assay (Thermo Fischer Scientific).

For nucleocytoplasmic fractionation of autopsy tissue, fractionation was performed with the NE-PER Nuclear and Cytoplasmic Extraction Kit #78833 (Thermo Fischer Scientific) according to the manufacturer’s protocol. For detection of proteins in the whole fraction, *Drosophila* larvae were solubilized in 8M urea. For the soluble and pelleted fraction, larvae were first solubilized in RIPA buffer as described above. The samples were spun down at 15000 rpm for 20 minutes and the soluble supernatant was set aside. Freshly prepared 8M urea buffer (Sigma) was added to the pellet and dissolved through vortexing. Samples were spun again at 15000rpm for 20 minutes and urea-soluble pellet fraction was collected. A small amount of sample buffer dye was added and urea-buffered protein samples were run immediately on SDS-PAGE without heating. For immunoblot, 10-50ug of total protein sample was mixed with 4x Laemmli buffer (Bio-Rad) and heated at 98 °C for 10 min. The protein samples were run on 4–15% SDS Mini-PROTEAN TGX Precast Gels (Bio-Rad) and transferred to nitrocellulose membrane. TBST (50 mM Tris-HCl pH 7.4, 1% Triton X-100) with 5% non-fat milk (Bio-Rad) was used for blocking. Primary antibodies were used as below: rabbit anti-Ref(2)p [a gift from Gabor Juhasz, Eotvos Lorand University, Budapest, Hungary] 1:1000; chicken anti-GFP (Abcam) 1:1000; guinea pig anti-CP1 [a gift from Patrick Dolph, Dartmouth College, NH, USA]; mouse anti-poly ubiquitin FK1 (Enzo Life Sciences) 1:1000; mouse anti-beta actin (clone C4) (EMD Millipore); rabbit anti-TFEB (Bethyl Laboratories) 1:2000, rabbit anti-histone H3 (Abcam) 1:1000.

### Electroretinogram (ERG) Assay

For ERG recordings, *Rh1-GAL4*/*UAS-LacZ* and *Rh1-GAL4*/*UAS-(G4C2)*_*30*_ flies were aged at 25°C in 12h light/12h dark cycle. ERG recordings were performed as described^66^. In brief, adult flies were immobilized on a glass slide by glue. A reference electrode was inserted in the thorax and a recording electrode was placed on the eye surface. Flies were maintained in the darkness for at least 2 minutes prior to 1-second flashes of white light pulses (LED source with daylight filter), during which retinal responses were recorded and analyzed using WinWCP (University of Strathclyde, Glasgow, Scotland) software. At least 5 flies were examined for each genotype and timepoint.

### Transmission Electron Microscopy (TEM)

*rh1-GAL4/UAS-lacZ* and *rh1-GAL4/UAS-(G4C2)*_*30*_ flies were aged at 25°C in 12h light/12h dark cycle. Retinae of adult flies were processed for TEM imaging as previously described (Chouhan et al., 2016). 3 flies were examined for each genotype and timepoint.

### Plasmids Source and Construction

pSF-CAG-Amp (0G504) were purchased from Oxford Genetics. We generated a mammalian expressing plasmid pSF-(G4C2)47-VFH (V5-Flag-His), which can express 47 HRE repeats with 3 different tags to monitor expression of DPRs (polyGA, polyGP, and polyGR). pEGFP-(GA,GR, or PR)50 was obtained from Davide Trotti (Wen et. al 2014) and the GFP cDNA sequence was replaced with mCherry by digesting with BamHI and XhoI.

### TFEB-GFP Hela cell culture and transfection

Hela cell line with stable expressing TFEB::GFP was a gift from Dr. Shawn Ferguson at Yale University. Hela cells were grown in DMEM media (Invitrogen) supplemented with 10% fetal bovine serum (Hyclone Laboratories Inc.) as previous described^93^. Transfection was performed using Lipofectamine 2000 (Invitrogen) according to the manufacturer’s instructions. Briefly, 1-2 µg of cDNA was diluted into 100 µl of Opti-MEM I Medium (Invitrogen) and mixed gently. Lipofectamine 2000 mixture was prepared by diluting 2-4 µl of Lipofectamine 2000 in 100 µl of Opti-MEM I Medium. The ratio of DNA to Lipofectamine 2000 used for transfection was 1:2 as indicated in the manual. The DNA-Lipofectamine 2000 mixture was mixed gently and incubated for 20 min at room temperature. Cells were directly added to the 200 µl of DNA-Lipofectamine 2000 mixture. After 48 h, transfected Hela cells were treated with EBSS medium for 3 h for starvation.

### Immunofluorescence analysis of Hela cells

Hela cells were fixed with 4% PFA at room temperature for 15 min, washed three times with PBS, permeabilized for 10 min with 1% PBTX, washed another three times with PBS, and blocked for 1 hr at room temperature with 10% normal goat serum (Sigma) diluted in 0.1% PBTX. Cells were then incubated overnight at 4°C with primary antibody mouse anti-Flag antibody (1:1000, Sigma). After three washes in PBS (5 min each), cells were incubated for 1 hr at room temperature with secondary antibodies (goat anti-Alexa Fluor 568, 1:250; Invitrogen) diluted in the blocking solution. Cells were washed three times in PBS and mounted with Prolong Gold anti-fade reagent with DAPI (Cell Signaling).

### Antibodies

For immunofluorescence experiments, the following antibodies were used: Rabbit anti-dsRed (Clontech #632496) 1:1,000; mouse anti-poly ubiquitin FK1 (Enzo Life Sciences #BML-PW8805) 1:200; rabbit anti-Ref2p^93^ (a gift from Gabor Juhasz, Eotvos Lorand University, Budapest, Hungary) 1:1000; guinea pig anti-Mitf^39^ (a gift from Francesca Pignoni, SUNY Upstate, New York, USA) 1:500; rat anti-HA (Sigma #11867423001) 1:200; mouse anti-Flag antibody (Sigma #F3165) 1:1000. For Western experiments: rabbit anti-Ref(2)p (from Gabor Juhasz), 1:1000; chicken anti-GFP (Abcam ab13970) 1:1000; guinea pig anti-CP1^56^ (a gift from Patrick Dolph, Dartmouth College, NH, USA); mouse anti-poly ubiquitin FK1 (Enzo Life Sciences #BML-PW8805) 1:1000; mouse anti-beta actin (clone C4) (EMD Millipore, MAB1501); rabbit anti-TFEB (Bethyl A303-673A) 1:2000; rabbit anti-histone H3 (Abcam) 1:1000.

### Collection of human autopsied tissue

Human autopsied tissue used for these data are described in detail in Supplementary Table 2. The use of human tissue and associated decedents’ demographic information was approved by the Johns Hopkins University Institutional Review Board and ethics committee (HIPAA Form 5 exemption, Application 11-02-10-01RD) and from the Ravitz Laboratory (UCSD) through the Target ALS Consortium.

### Statistics

All quantitative data were derived from independent experiments. Each n value representing biological replicates is indicated in the figure legends. Statistical tests were performed in Prism version 8.3.1 or Microsoft Excel 16.34 and were performed as marked in the figure legends. All statistical tests were two-sided. Results were deemed significant when the P value α = 0.05, (p) * = p<0.05, ** = p<0.01, *** = p<0.001, n.s. = not significant. No statistical methods were used to predetermine sample size. The investigators were not blinded during experiments.

### Reagent Table

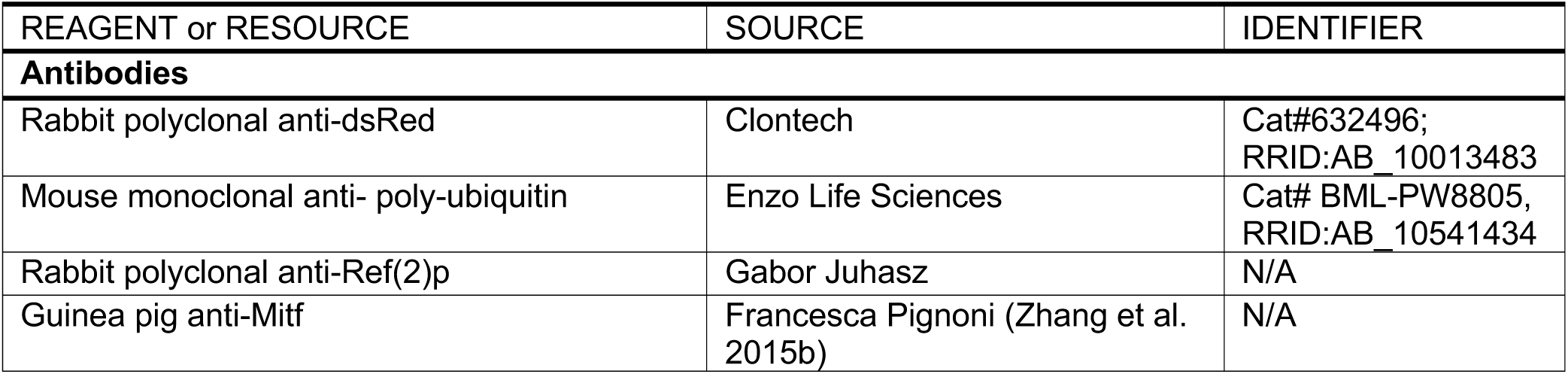
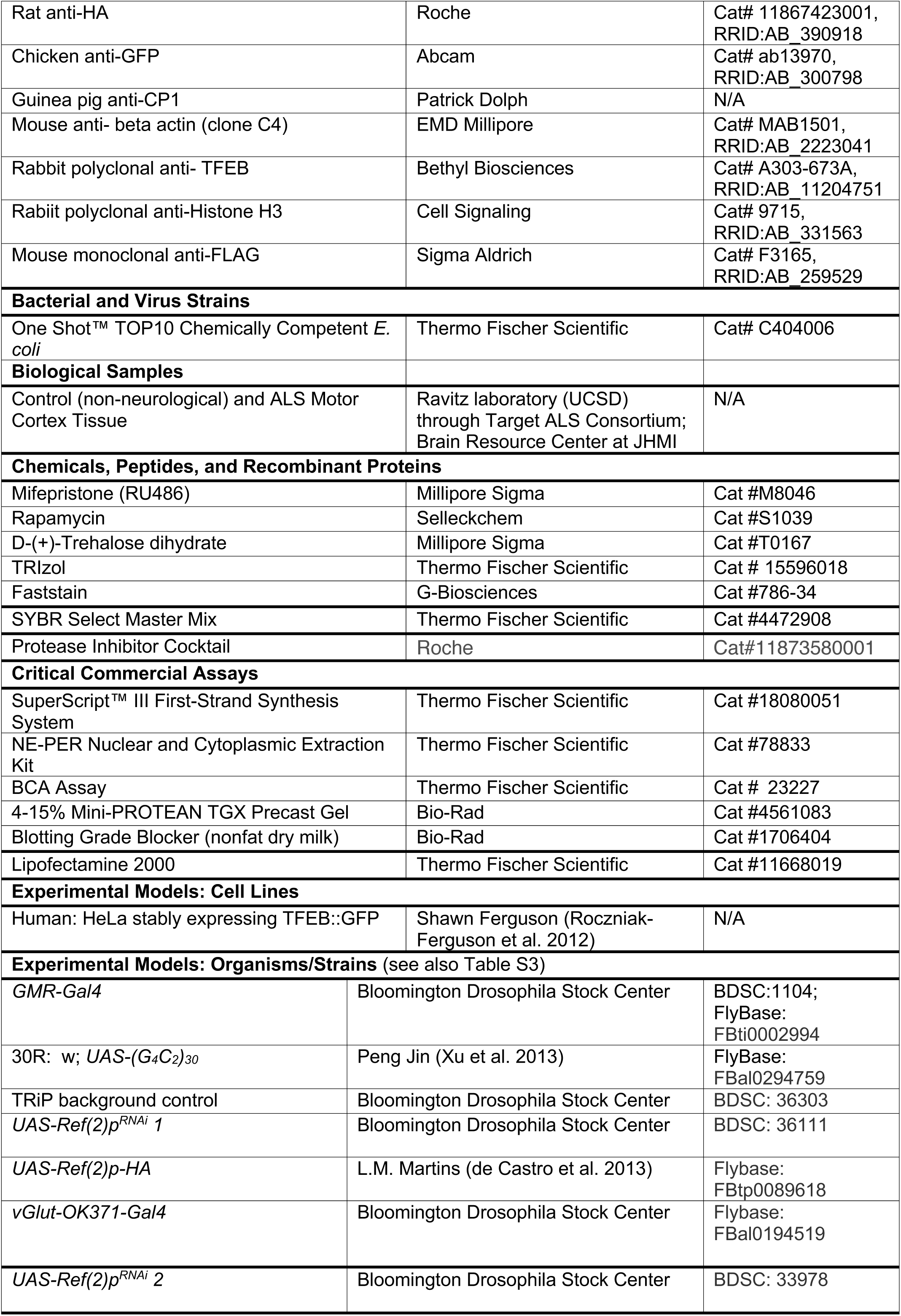
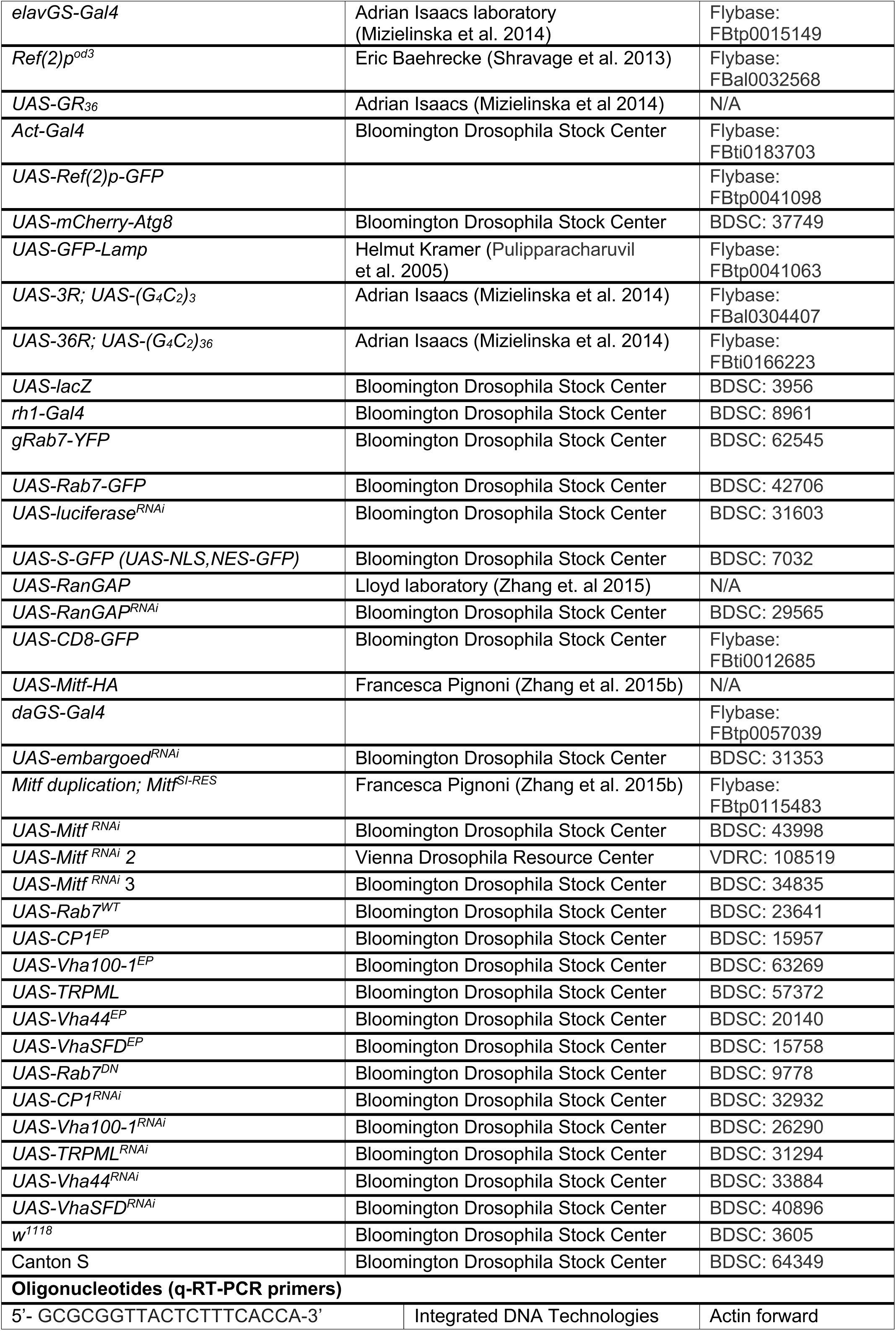
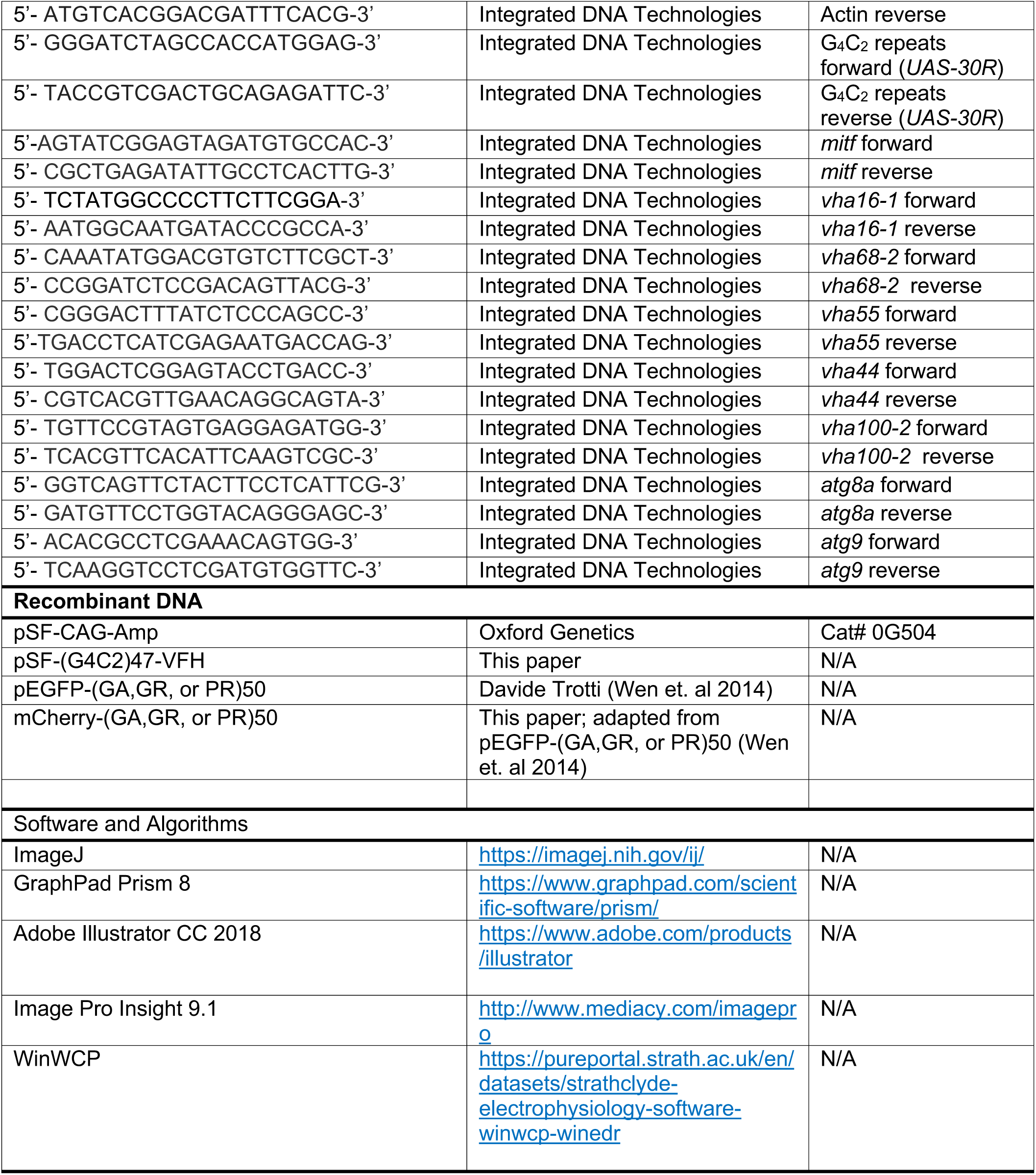

## Acknowledgments

This work was supported by NINDS R01NS082563 (T.E.L), R01NS094239 (T.E.L and J.D.R.), F31 NS100401 (K.M.C.), ALSA (T.E.L., K.Z. and J.D.R.), and Target ALS (T.E.L., K.Z., J.D.R., and H.J.B.). K.M.C is a recipient of the P.E.O Scholar Award. H.J.B. is an Investigator of the Howard Hughes Medical Institute. We thank Francesca Pignoni, Udai Pandey, Peng Jin, Adrian Isaacs, Eric Baehrecke, Helmut Kramer, Francesca Pignoni, Gábor Juhász, Patrick Dolph, L. Miguel Martins, the Bloomington *Drosophila* Stock Center (NIH P40ODO18537) and Vienna *Drosophila* Research Center for *Drosophila* lines and/or antibodies and Shawn Ferguson and Davide Trotti for cell lines and constructs. The Johns Hopkins NINDS Multiphoton Imaging Core (NS050274) provided imaging equipment and expertise.

## Author Contributions

K.M.C., K.Z., and T.E.L. conceived the study and designed experiments; K.M.C., K.Z. K.R., H.S., and K.M. performed *Drosophila* and cell culture experiments and analyzed data. J.G. (supervised by J.D.R.) and K.M.C. performed human tissue experiments. M.S. and Z.Z. performed the TEM experiments. K.M.C., K.Z., H.J.B., and T.E.L. wrote and edited the manuscript. T.E.L. supervised the project and acquired funding.

---

**Supplemental Figure 1 (Related to Figure 1):**
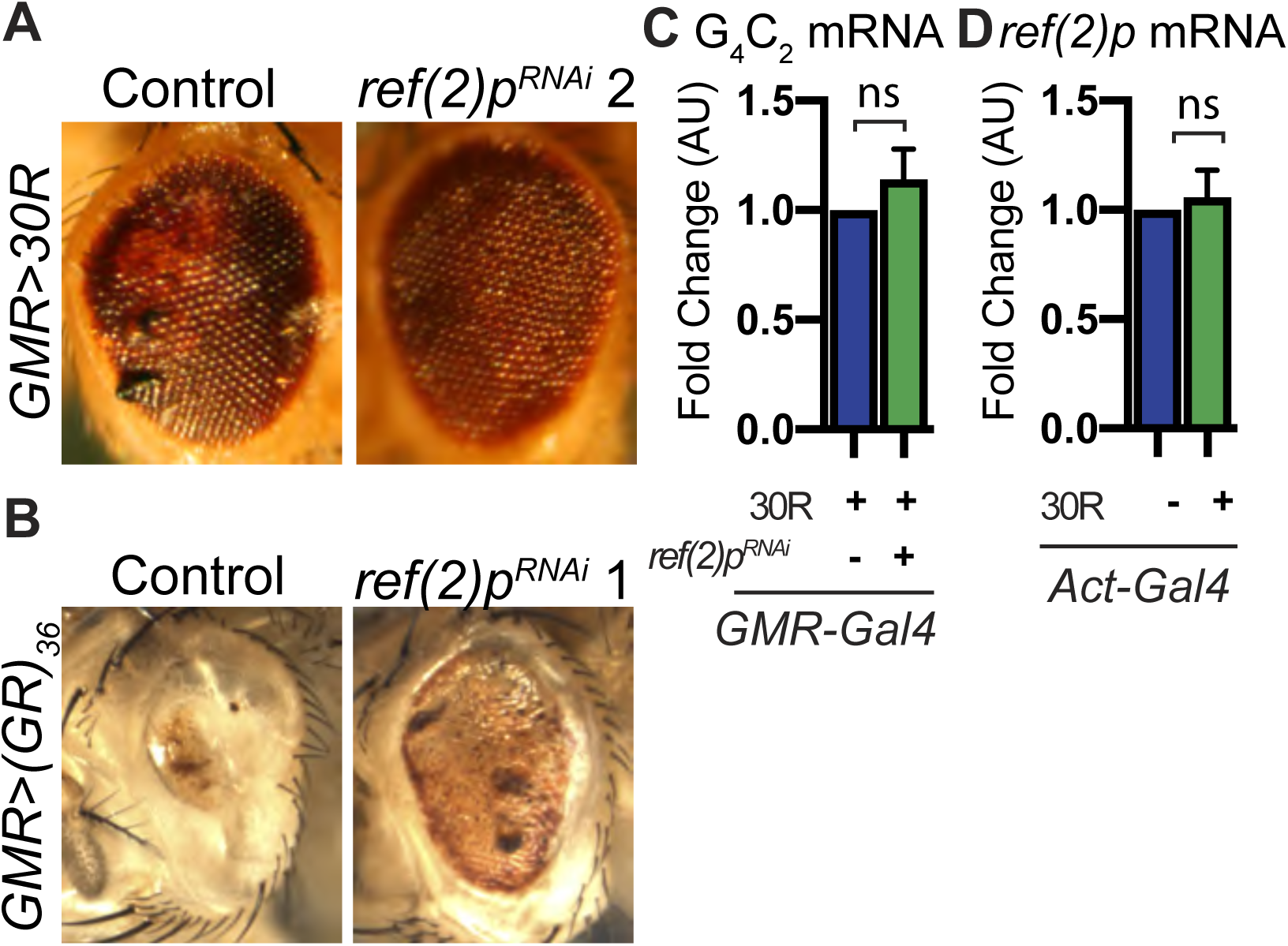
Ref(2)p/p62 genetically modifies G4C2-HRE. A) Control and a second *Ref(2)p*^*RNAi*^ line (*Ref(2)p*^*HMS00938*^, see methods for detailed line information) in *GMR*>30R expressing *Drosophila* eyes at 15 days of age. B) *GMR-Gal4* expressing an alternate codon *poly-GR*_*36*_ alone or co-expressed with *Ref(2)p*^*RNAi*^ *(Ref(2)p*^*HMS00551*^) at 15 days of age. D) Levels of G4C2 RNA measured by qPCR (See methods) in *Drosophila* expressing 30R under the control of *GMR-Gal4* either with or without *Ref(2)p RNAi*, n=3, Student’s t-test. E) Levels of *Ref(2)p* RNA in *Drosophila* larva expressing 30R ubiquitously under control of the *Actin-Gal4*, n=5, Student’s t-test. Data represented are mean ± SEM. n.s., not significant.

**Supplemental Figure 2 (Related to Figure 2):**
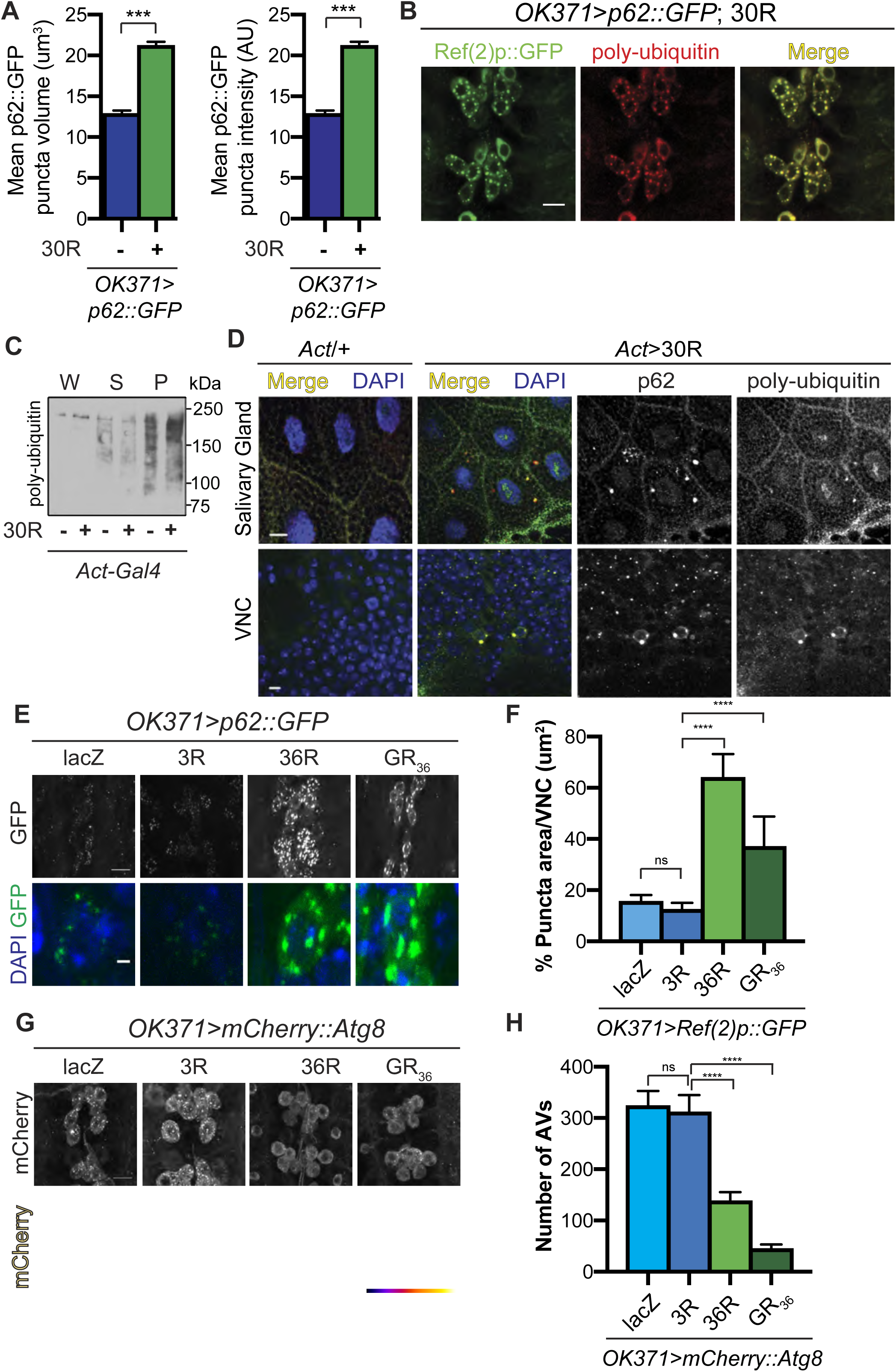
p62::GFP aggregates in C9-ALS fly models and co-localizes with poly-ubiquitin. A) Quantification of mean intensity and mean volume of p62::GFP puncta in *OK371/*+ or *OK371*>30R motor neuron cell bodies, n=5 Student’s t-test with Welch’s correction ***p<.0001. B) Example image of *Drosophila OK371*-Gal4 motor neurons expressing 30R; *p62::GFP* (green) co-stained with an anti-poly-ubiquitin (red). C) Western blot of anti-poly-ubiquitin showing the whole (W), supernatant (S) and pellet (P) fractions of *Drosophila* L3 larvae ubiquitously expressing 30R under the control of *Act-Gal4*. D) Images of *Drosophila* L3 larvae *Act-Gal4*/+ or *Act*>30R ubiquitously expressing G4C2 repeats showing anti-Ref(2)p (red) and anti-ubiquitin (green) staining in the larval salivary gland (top), ventral nerve cord (VNC) (bottom). E) *OK371*>*p62::GFP* showing the VNC (top) or a representative cell body (bottom) with two control (*UAS-lacZ* or *UAS-3R*) and two alternate C9-ALS model lines, one expressing a G4C2 repeat expansion (*UAS-36R*) and another expressing the alternate codon arginine dipeptide GR (*UAS-GR*_*36*_). F) Quantification of the total GFP+ area of p62::GFP positive puncta in A). G) *OK371*>*mCherry::Atg8* showing the VNC (top) or a representative cell body (bottom, scale bar represents 1um) with two control overexpression (*UAS-lacZ* or *UAS-3R*) and two alternate C9-ALS model lines, one expressing a G4C2 repeat expansion (*UAS-36R*) and another expressing the alternate codon arginine dipeptide GR (*UAS-GR*_*36*_). H) Quantification of mCherry:Atg8 puncta in the motor neuron cell bodies in (G). Data represented are mean ± SEM. Scale bar represents 10µm. n.s., not significant, ***p<0.001, ****p<0.0001.

**Supplemental Figure 3 (Related to Figure 3):**
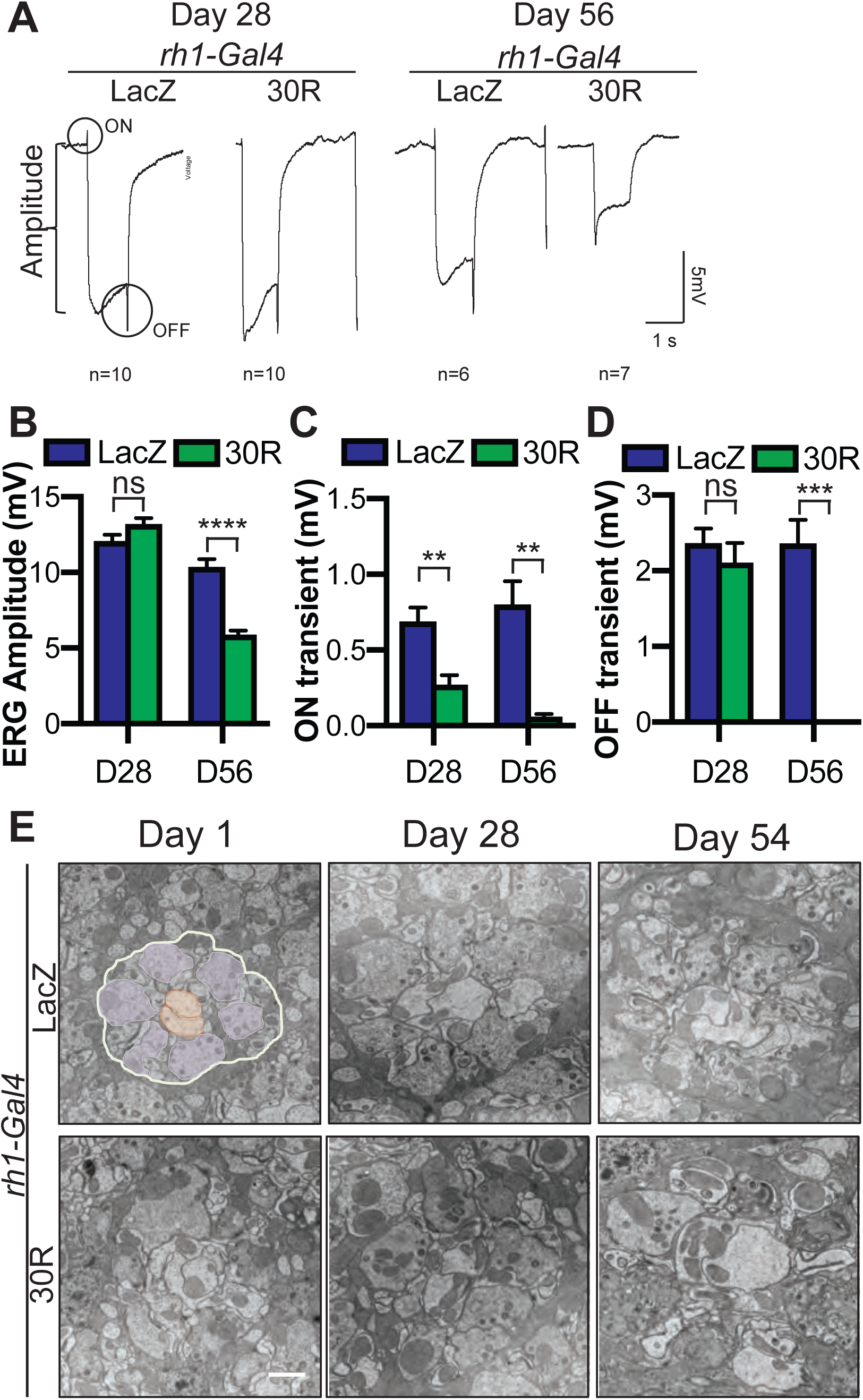
Progressive synapse degeneration in G4C2-expressing photoreceptor neurons. A) Representative electroretinogram (ERG) of *rhodopsin1-Gal4* driving *LacZ* (control) or 30R in adult eyes at Day 28 or Day 56 after eclosion. B) Quantification of mean ERG amplitude in A. C) Quantification of mean ON transient amplitude in A. D) Quantification of mean OFF transient amplitude in A. Student’s t-test, n = 10, 10, 6, 7. E) Transmission electron micrscopy (TEM) of terminals (synapses) in *Rhodopsin1-Gal4* driving *UAS-LacZ* (control) or 30R adult eyes at Day 1, Day 28, and Day 54 after eclosion. For the Day 1 control image, presynaptic terminals are pseudocolored in purple and post-synaptic terminals are pseudocolored in orange. Data represented are mean ± SEM. Scale bar represents 1µm. n.s., not significant *, p<0.05, **, p<0.01, ***p<0.001, ****p<0.0001

**Supplemental Figure 4 (Related to Figure 2 and 3):**
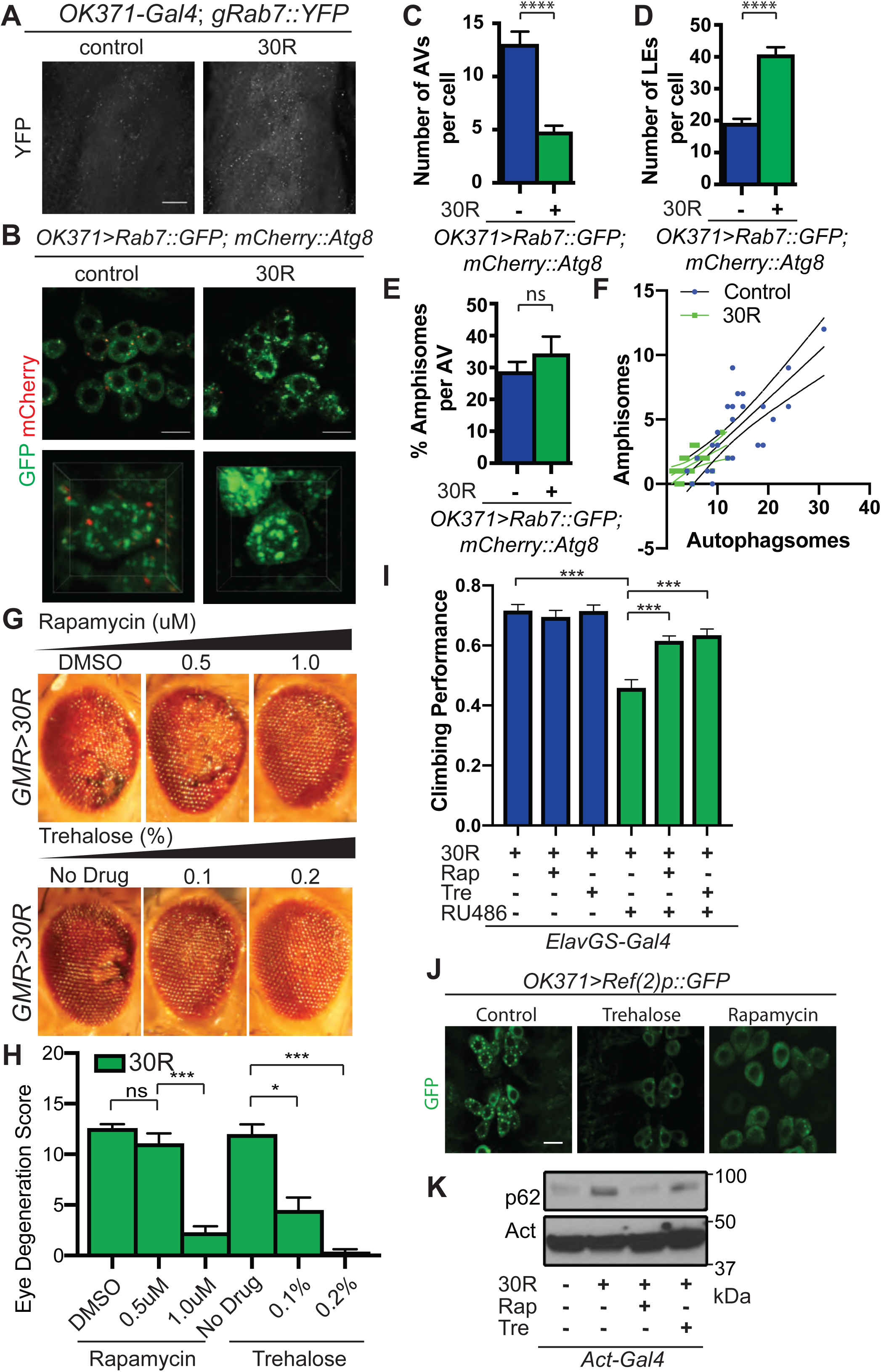
Rescuing G4C2-mediated lysosome defects reduces neurodegeneration. A) *OK371*/+ or *OK371*>30R with genomically tagged *Rab7::YFP* in the larval ventral nerve cord. B) Control *OK371* motor neurons co-expressing *UAS-Rab7::GFP* and *UAS-mCherry::Atg8* in the ventral nerve cord (top). Representative single cell (bottom). C) Quantification of number of Atg8+ vesicles autophagic vesicles per cell, Student’s t-test. D) Quantification of number of Rab7+ late endo/lysosomes (LE) per cell (Student’s t-test). E) Mean number of amphisomes (co-localized mCherry::Atg8 and Rab7::GFP) per cell normalized to the total number of autophagosomes. F) Number of amphisomes (co-localized mCherry::Atg8 and Rab7::GFP) spots plotted against number of autophagic vesicles for *OK371*/+ control and *OK371*>30R motor neurons. G) 15-day old *Drosophila* eyes expressing 30R using *GMR-Gal4* of flies fed with increasing concentrations of rapamycin (DMSO, 0.5 µM, or 1µM) and trehalose (no drug, 0.1%, or 0.2%). H) Quantification of degeneration in (G), One-way ANOVA, followed by multiple comparisons. I) Adult flies expressing 30R under the control of the inducible, pan-neuronal elavGS-Gal4 driver have decreased climbing ability after 7 days compared to non-induced controls, which is rescued by supplementing the food with rapamycin or trehalose, One-way ANOVA with Bonferroni’s Multiple Comparison’s test. (J) *Drosophila* motor neurons expressing *Ref(2)p::GFP* and 30R from 3^rd^ instar larvae fed DMSO (control) or 1.0µM rapamycin or 0.2% trehalose. (E) Western of whole *Drosophila* larvae expressing no repeats or 30R and fed DMSO or 1.0uM rapamycin or 0.2% trehalose and blotted for Ref(2)p. Data represented are mean + SEM. Scale bars represent 10µm. n.s., not significant *, p<0.05, ***,p<0.001, ****,p<0.0001

**Supplemental Figure 5 (Related to Figure 4).**
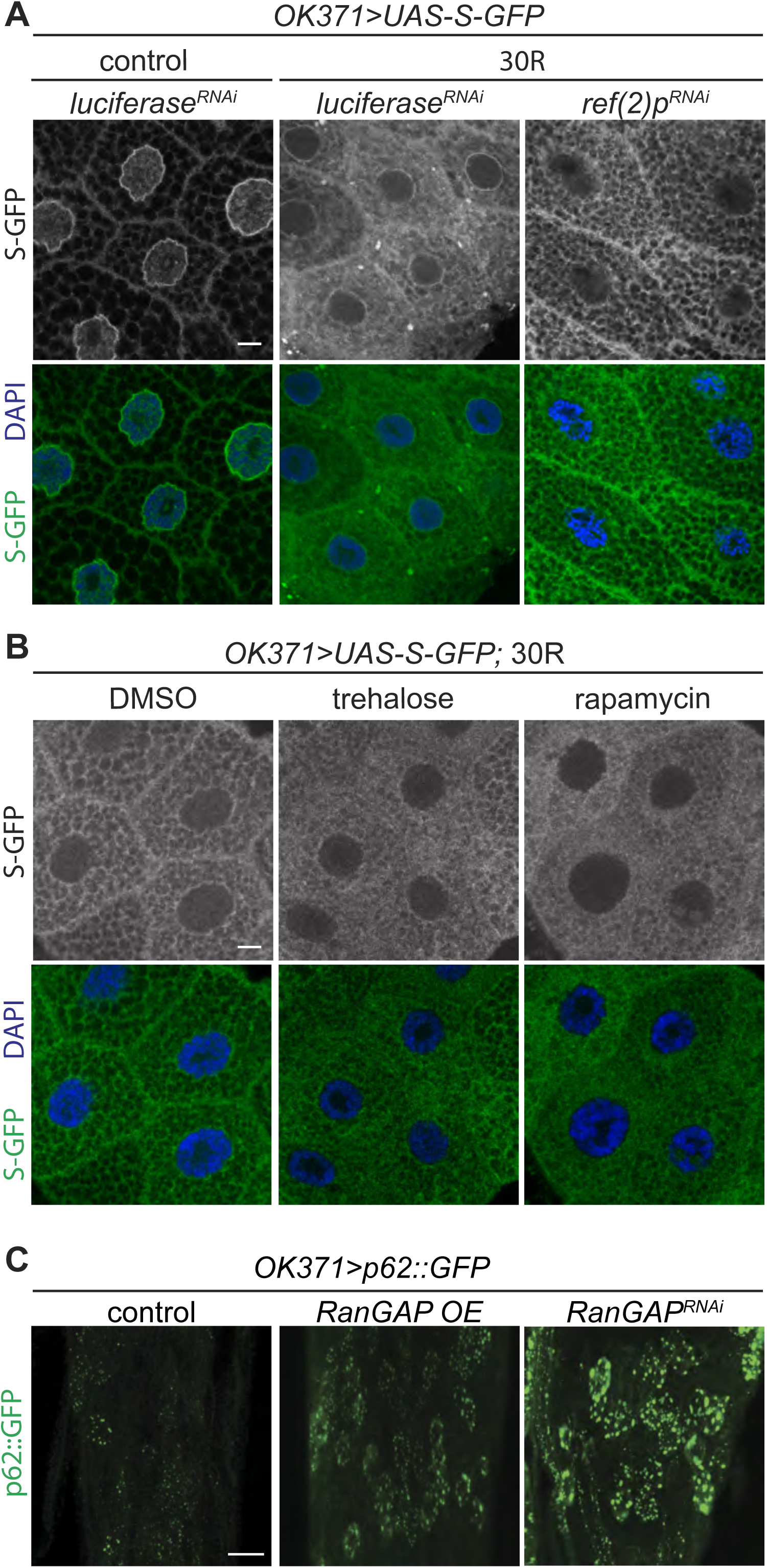
Nucleocytoplasmic transport disruption is upstream of autophagic defects. (A) Images of the nucleocytoplasmic transport marker shuttle-GFP (S-GFP) in control and *OK371*>*30R* with either *luciferase* (control) or *Ref(2)p* RNAi expressed in *Drosophila* salivary glands. (B) Images of L3 salivary gland expressing S-GFP in *OK371*>*30R Drosophila* after feeding supplemented with either control (DMSO), 0.2% trehalose, or 1.0µm rapamycin. C) Representative images of ventral nerve cords expressing either control, knockdown of *RanGAP* (*UAS-RanGAP*^*RNAi*^), of overexpression of RanGAP (*UAS-RanGAP)*, a master regulator of nucleocytoplasmic transport along with autophagy receptor Ref(2)P::GFP in *OK371* motor neurons. Data represented are mean ± SEM. Scale bars represent 10µm.

**Supplemental Figure 6 (Related to Figure 6).**
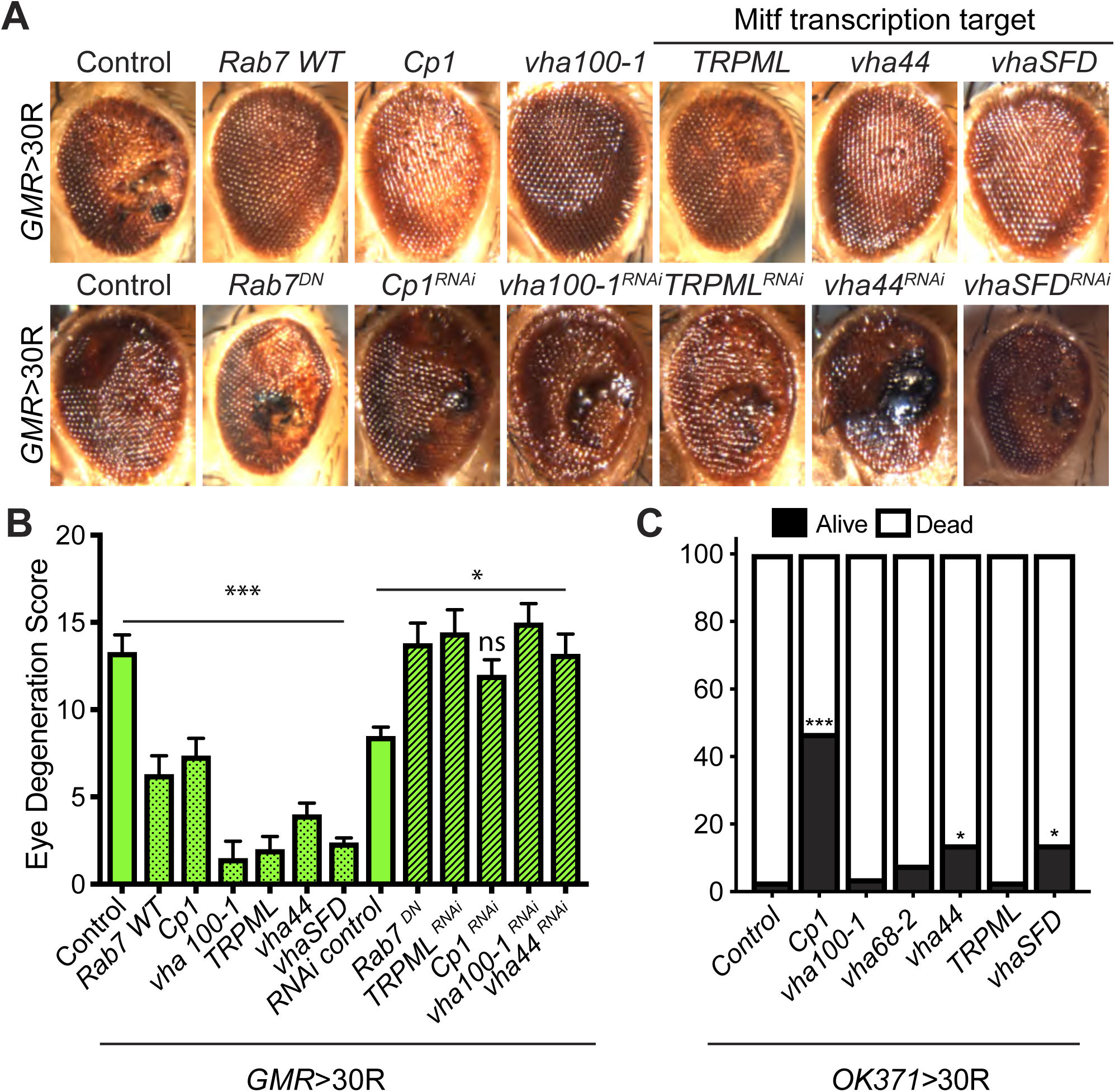
Genetic manipulation of lysosomes rescues degeneration caused by G4C2 expression. A) 15-day old *Drosophila* eyes expressing 30R under the control of the *GMR-Gal4* driver accompanying overexpression (top) or knockdown (bottom) of lysosomal genes. (B) Quantification of external eye degeneration in (A), see methods. One-way ANOVA, *,p<0.05, ***,p<0.001 N = 10, 10, 10, 10, 8, 4, 5, 8 adult flies. (C) Percent of pupal eclosion in *Drosophila* expressing 30R under the control of the motor neuron *OK371-Gal4* driver with *UAS-lacZ* (control), or overexpression of lysosomal genes N = 89, 120, 88, 146, 119, 123, 105 pupa from 3 replicates, Fisher’s exact test, *,p<.0.05, ***,p<0.001 Data represented are mean ± SEM.

**Supplemental Figure 7 (Related to Figure 7).**
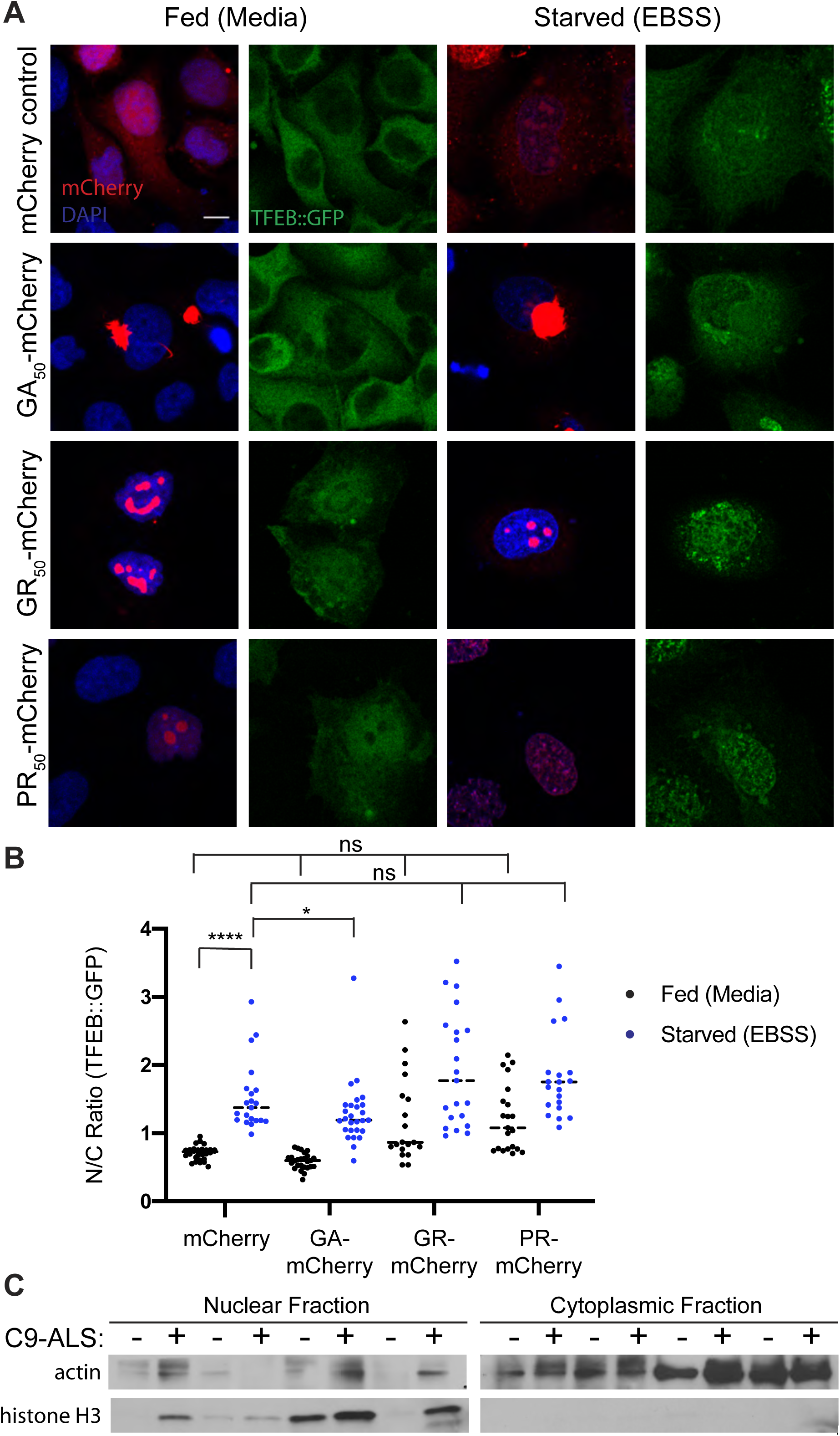
DPRs affect TFEB import in HeLa Cells. A) Representative images of transfection of codon-optimized poly-GA-mCherry, poly-GR-mCherry, and poly-PR-mCherry into HeLa cells with stably incorporated TFEB-GFP. B) Quantification of (A), One-way ANOVA with multiple comparisons. C) Nuclear and cytoplasmic fractions of human motor cortex samples in Figure 7c blotted for cytoplasmic (actin) and nuclear (histone H3) housekeeping controls. Scale bars represent 20um. n.s., not significant *, p<0.05, **, p<0.01, ***p<0.001, ****p<0.0001

## Supplemental Tables

**Supplemental Table 1.** Candidate Screen of autophagy-related genes. Flies expressing 30 G4C2 repeats in the eye under control of GMR-GAL4 were crossed to the indicated UAS line and scored for enhancement (<0) or suppression (>0) as described (Zhang et al., 2015a).

| Drosophila stock | Effect on gene | Eye Score |
| --- | --- | --- |
| *UAS-Atg6<sup>RNAi</sup>* | RNAi line | -3.5 |
| *UAS-Atg18a<sup>RNAi</sup>* | RNAi line | -3 |
| *UAS-Atg1* | cDNA overexpression | -3 |
| *UAS-Atg7* | cDNA overexpression | -2.5 |
| *UAS-Atg101<sup>RNAi</sup>* | RNAi line | -2 |
| *UAS-Atg8a<sup>RNAi</sup>* | RNAi line | -2 |
| *UAS-Atg5<sup>RNAi</sup>* | RNAi line | -2 |
| *UAS-Atg6<sup>RNAi</sup>* | RNAi line | -2 |
| *UAS-Atg8a<sup>RNAi</sup>* | RNAi line | -2 |
| *UAS-Ref(2)<sup>pRNAi</sup>* | RNAi line | -2 |
| *UAS-Atg14<sup>RNAi</sup>* | RNAi line | -2 |
| *UAS-Atg16<sup>RNAi</sup>* | RNAi line | -1.5 |
| *UAS-Atg16<sup>RNAi</sup>* | RNAi line | -1.5 |
| *UAS-Atg17<sup>RNAi</sup>* | RNAi line | -1.5 |
| *UAS-Atg6<sup>RNAi</sup>* | RNAi line | -1 |
| *UAS-Atg8b<sup>RNAi</sup>* | RNAi line | -1 |
| *UAS-bch<sup>SRNAi</sup>* | RNAi line | -1 |
| *UAS-Atg8b<sup>RNAi</sup>* | RNAi line | -1 |
| *UAS-Atg9<sup>RNAi</sup>* | RNAi line | -0.5 |
| *UAS-Atg18a<sup>RNAi</sup>* | RNAi line | -0.5 |
| *UAS-Gyf<sup>RNAi</sup>* | RNAi line | -0.5 |
| *Atg6<sup>00096</sup>* | P-element insertion | 0 |
| *UAS-Atg4b<sup>RNAi</sup>* | RNAi line | 0 |
| *UAS-Atg17<sup>EP</sup>* | EP overexpression | 0 |
| *bch<sup>S58</sup>* | EMS mutagenesis | 0 |
| *UAS-Atg4a<sup>RNAi</sup>* | RNAi line | 0 |
| *UAS-Atg4a<sup>RNAi</sup>* | RNAi line | 0 |
| *UAS-Atg10<sup>RNAi</sup>* | RNAi line | 0 |
| *UAS-Atg16<sup>RNAi</sup>* | RNAi line | 0 |
| *UAS-Atg9<sup>RNAi</sup>* | RNAi line | 0 |
| *UAS-It<sup>RNAi</sup>* | RNAi line | 0 |
| *UAS-Atg7<sup>RNAi</sup>* | RNAi line | 0 |
| *Atg7<sup>d06996</sup>* | P-element insertion | 0 |
| *UAS-Atg4a<sup>RNAi</sup>* | RNAi line | 1 |
| *UAS-Atg8a<sup>RNAi</sup>* | RNAi line | 1 |
| *Bch<sup>S17</sup>* | EMS mutagenesis | 1 |
| *UAS-Atg8a<sup>EP</sup>* | EP overexpression | 1 |
| *UAS-Atg2<sup>EP</sup>* | EP overexpression | 1 |
| *UAS-Atg7<sup>RNAi</sup>* | RNAi line | 1 |
| *UAS-Atg13<sup>RNAi</sup>* | RNAi line | 1.5 |
| *UAS-Atg14<sup>RNAi</sup>* | RNAi line | 1.5 |
| *UAS-Atg3<sup>EP</sup>* | EP overexpression | 2 |
| *UAS-Atg18b<sup>RNAi</sup>* | RNAi line | 2 |
| *UAS-Atg2<sup>RNAi</sup>* | RNAi line | 2 |
| *Snap29<sup>B6-21</sup>* | EMS | 2 |
| *UAS-Atg3<sup>RNAi</sup>* | RNAi line | 3 |
| *UAS-Atg2<sup>RNAi</sup>* | RNAi line | 3 |
| *UAS-Atg18b<sup>RNAi</sup>* | RNAi line | 3 |
| *Atg4b<sup>P0997</sup>* | P-element insertion | 4 |
| *UAS-bchs-HA* | cDNA overexpression | 4 |

**Supplemental Table 2 (Related to Figure 7):**
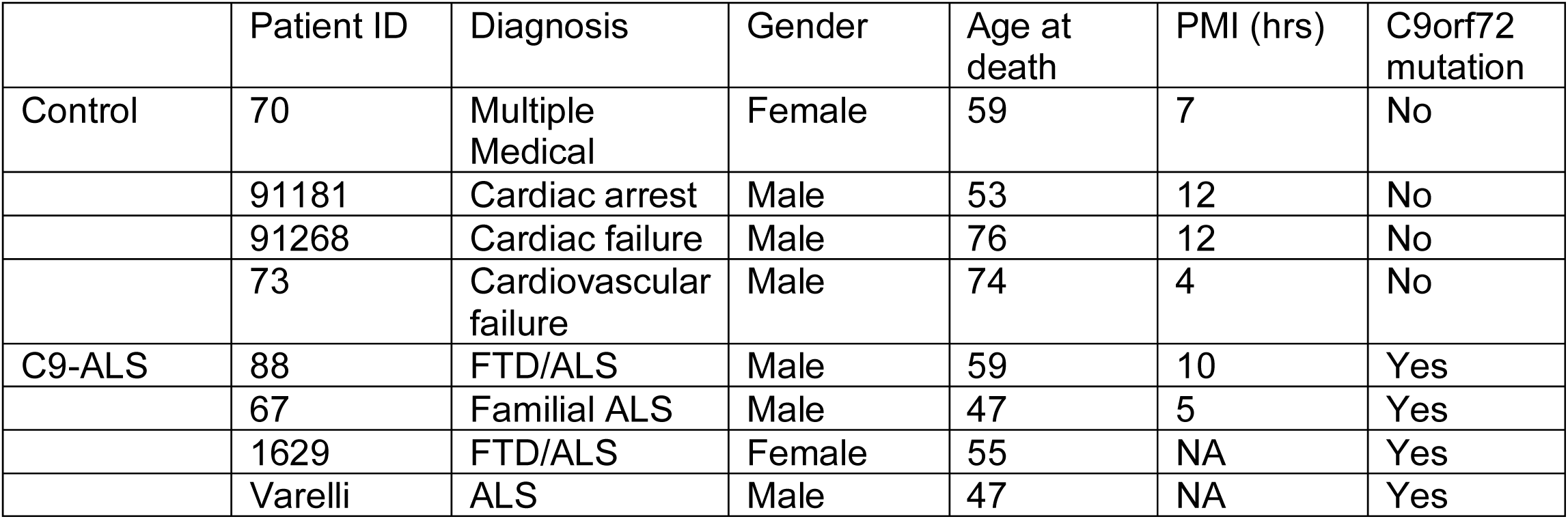
Demographics of human patients.

|  | Patient ID | Diagnosis | Gender | Age at death | PMI (hrs) | C9orf72 mutation |
| --- | --- | --- | --- | --- | --- | --- |
| Control | 70 | Multiple Medical | Female | 59 | 7 | No |
|  | 91181 | Cardiac arrest | Male | 53 | 12 | No |
|  | 91268 | Cardiac failure | Male | 76 | 12 | No |
|  | 73 | Cardiovascular failure | Male | 74 | 4 | No |
| C9-ALS | 88 | FTD/ALS | Male | 59 | 10 | Yes |
|  | 67 | Familial ALS | Male | 47 | 5 | Yes |
|  | 1629 | FTD/ALS | Female | 55 | NA | Yes |
|  | Varelli | ALS | Male | 47 | NA | Yes |

**Supplemental Table 3.**
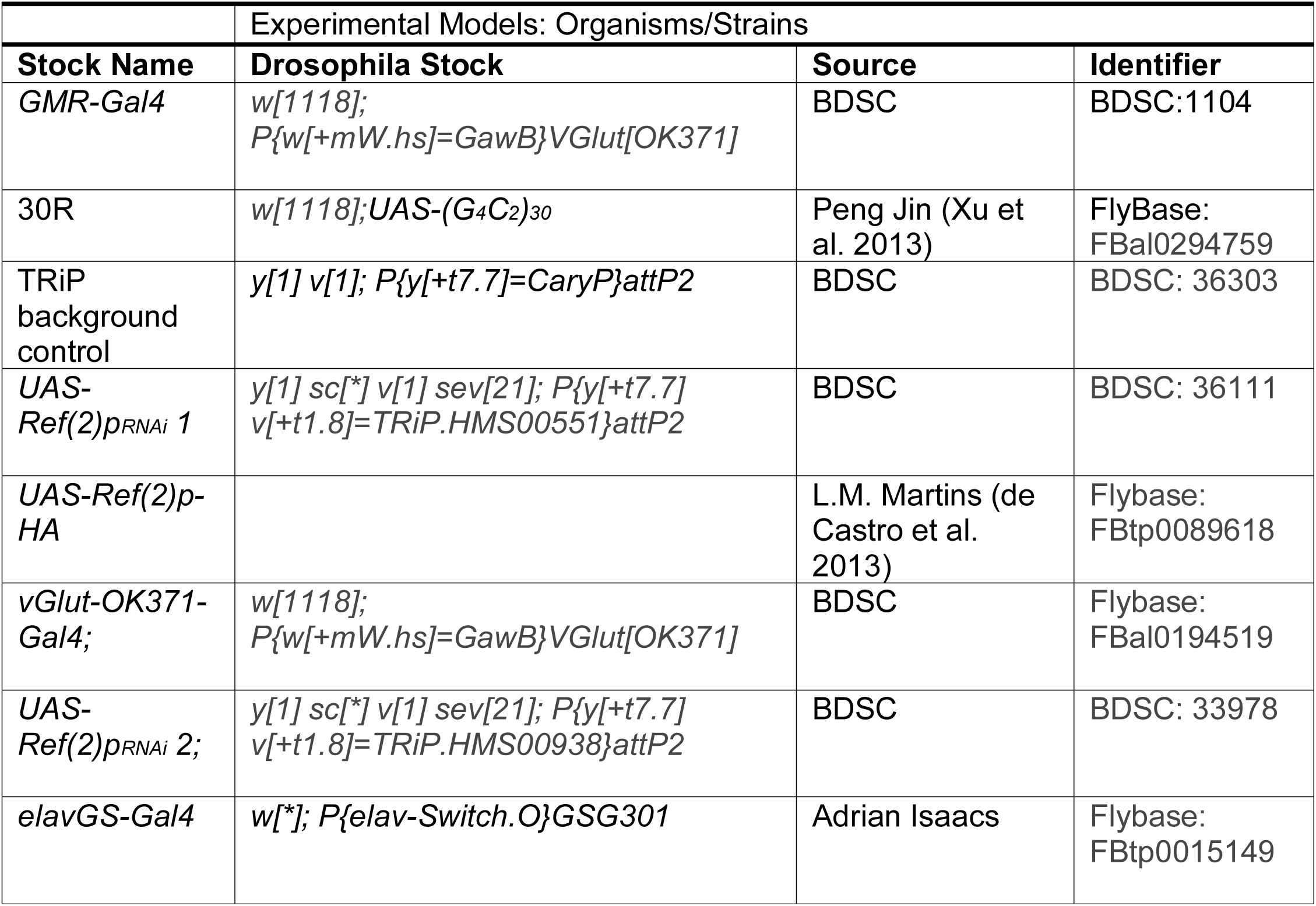
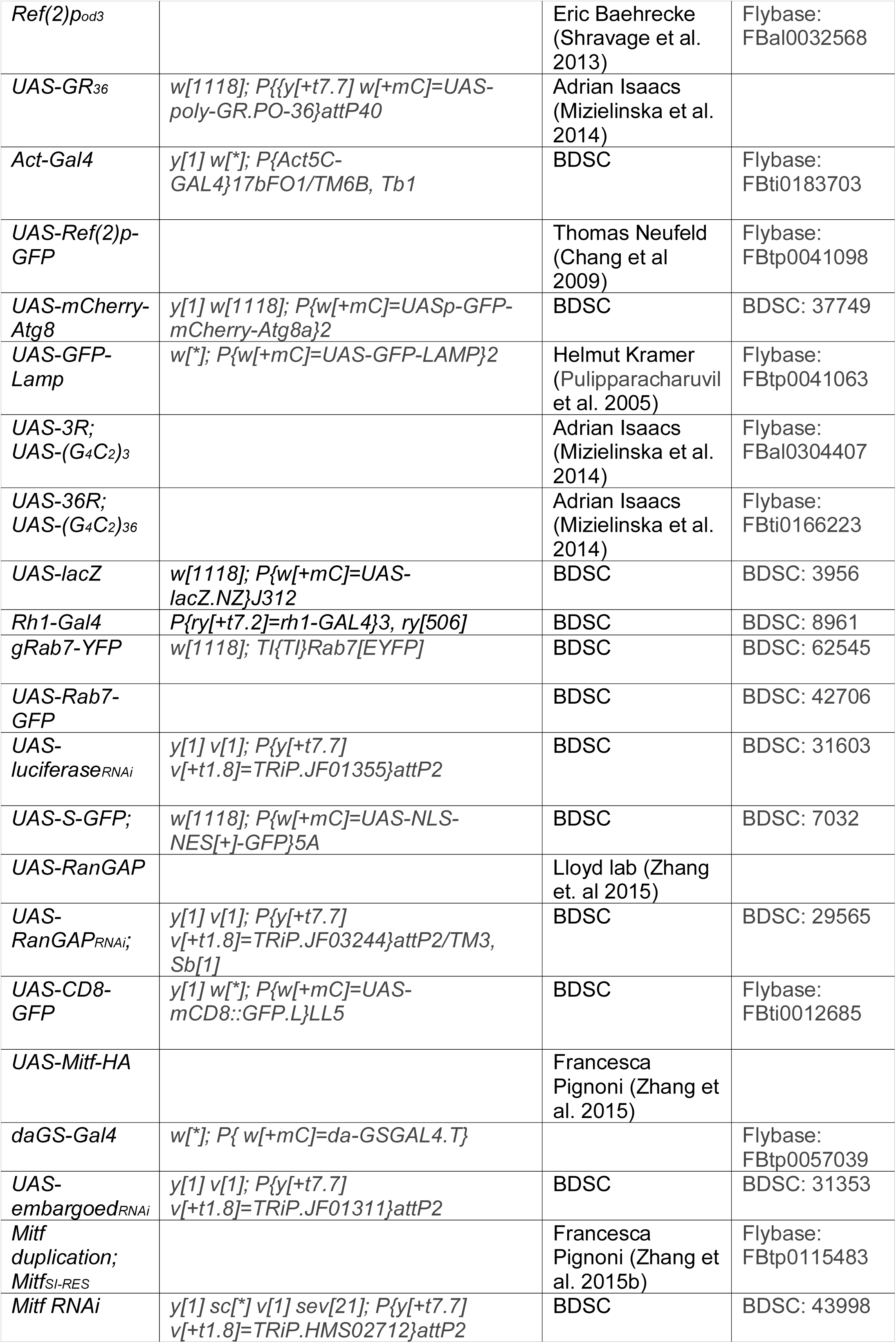
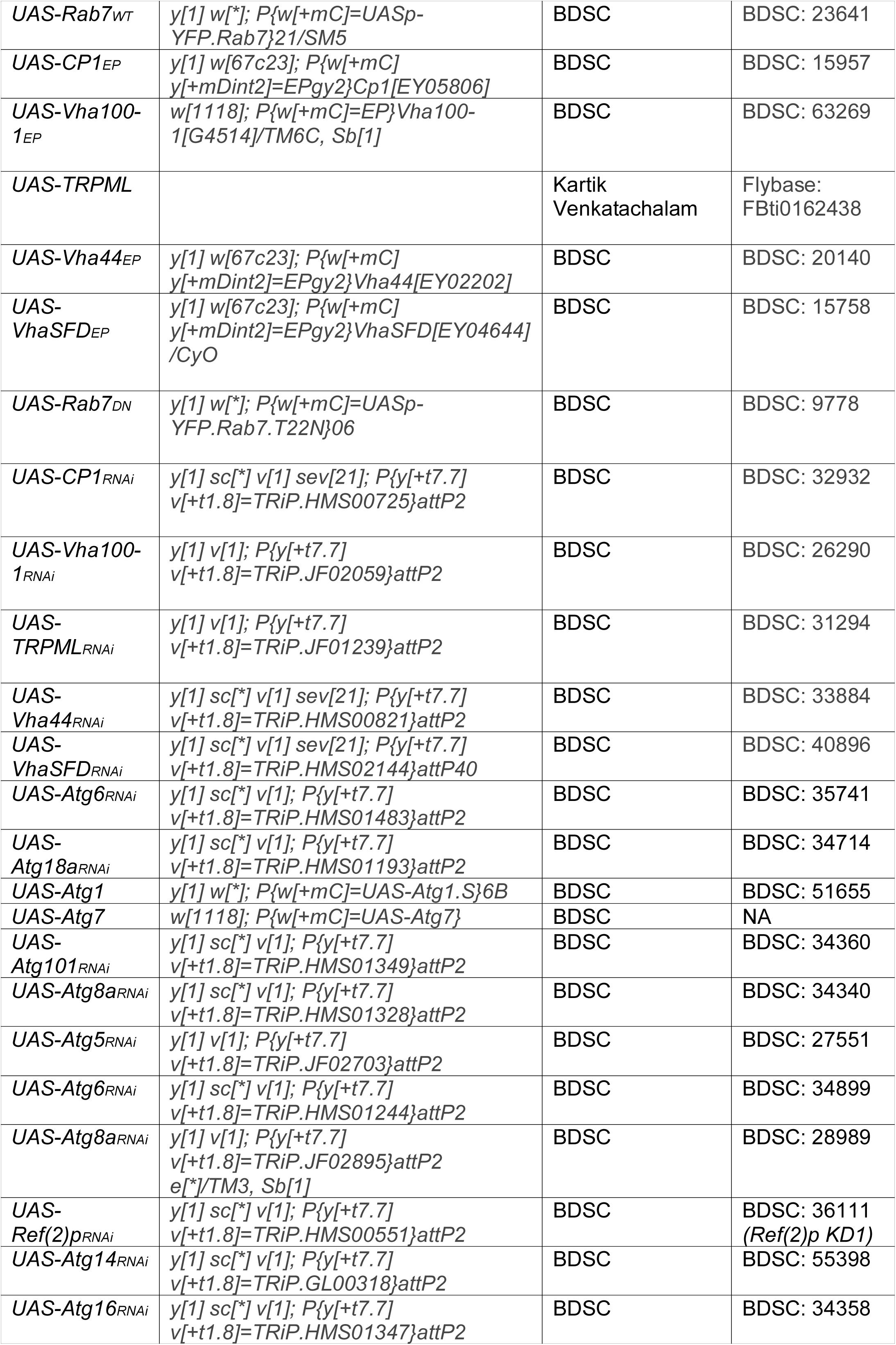
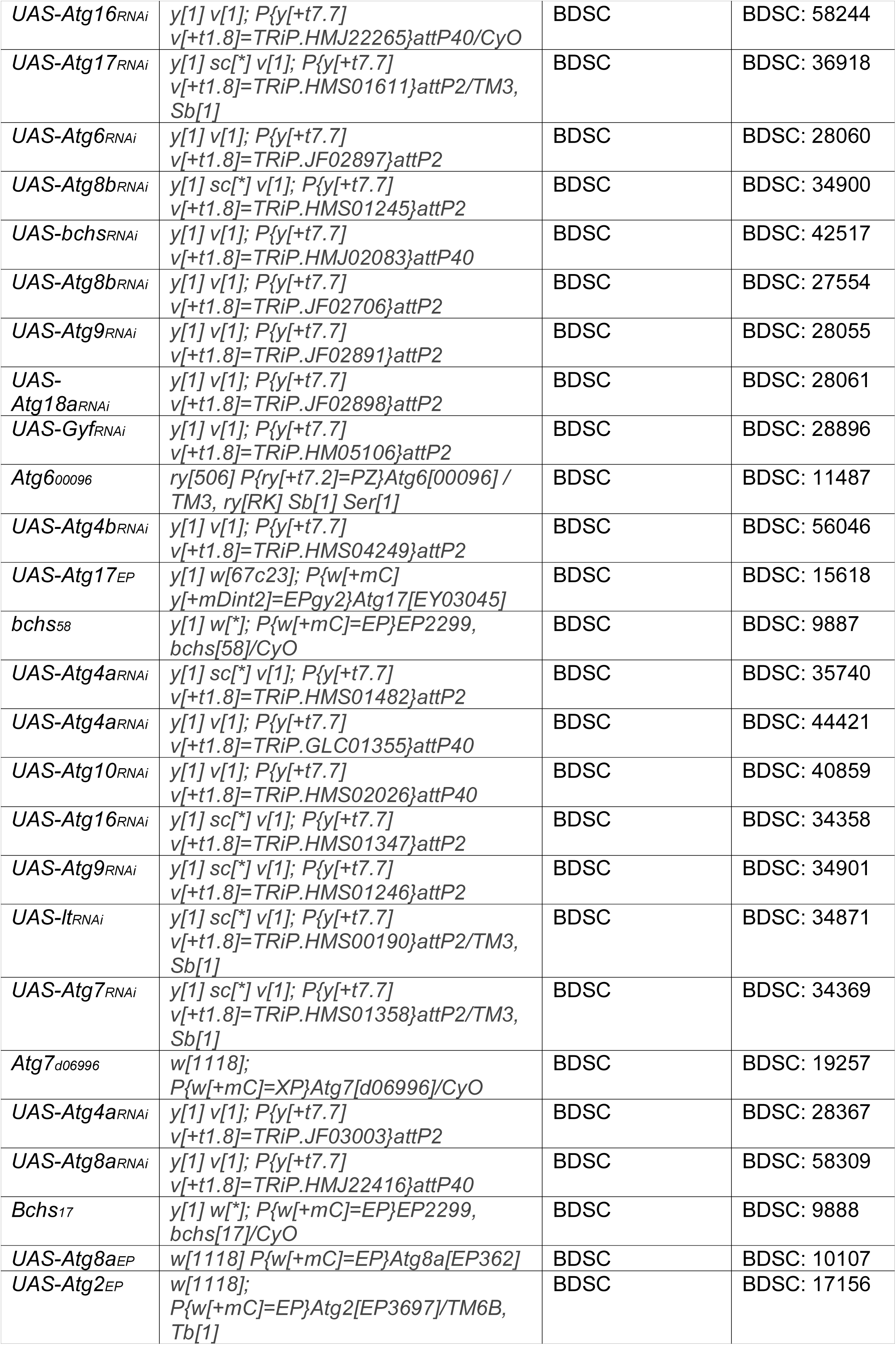
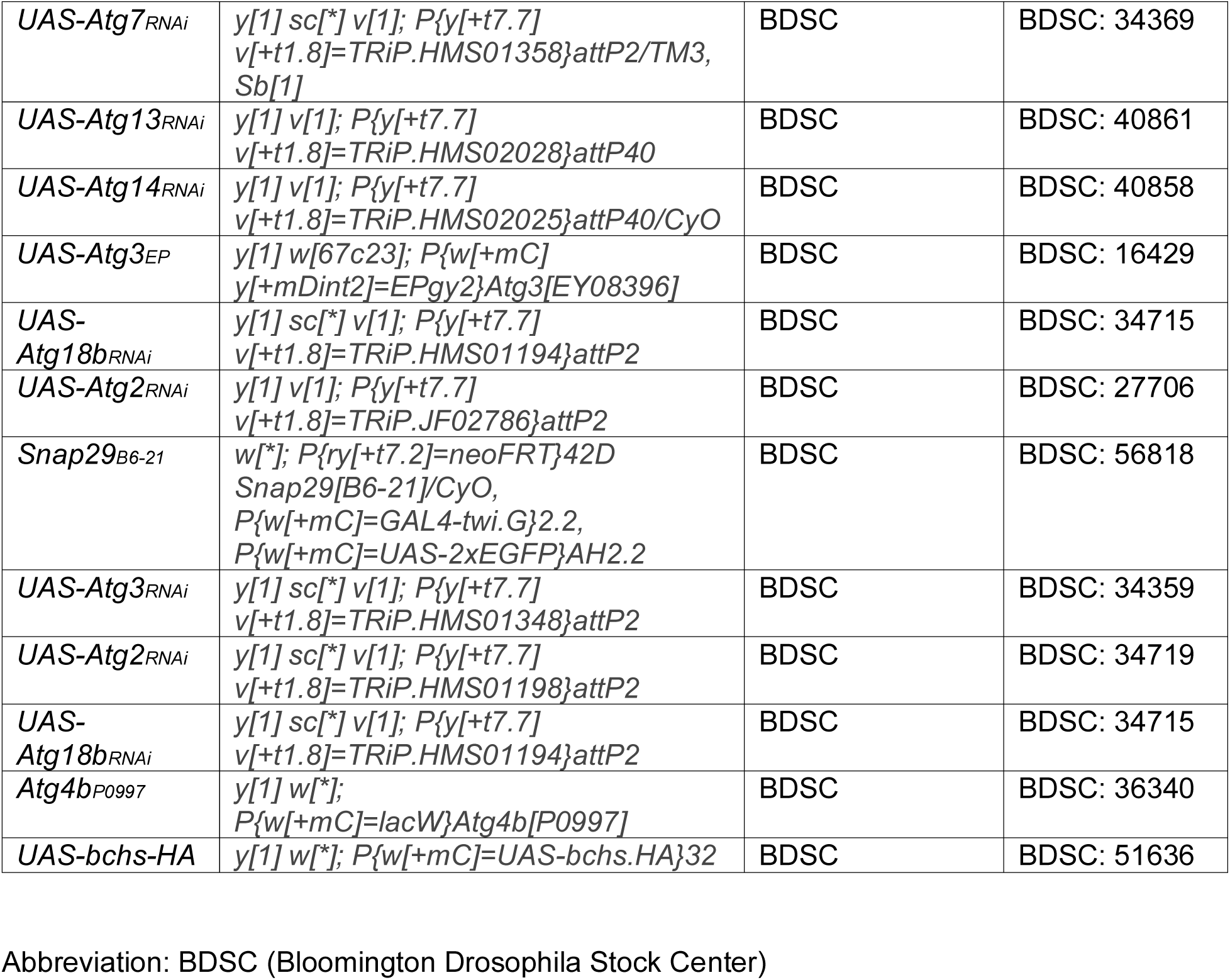
*Drosophila* Stocks used in this study.

